# Evolutionary Remodelling of a Remnant GET-Pathway Factor into PEX38, a Novel and Essential Peroxin in *Euglenozoa*

**DOI:** 10.1101/2025.11.03.686288

**Authors:** Chethan K. Krishna, Stefan Gaussmann, Hirak Das, Martin Jung, Silke Oeljeklaus, Michael Sattler, Bettina Warscheid, Vishal C. Kalel, Ralf Erdmann

## Abstract

PEX19 is a cytosolic receptor that directs membrane proteins post-translationally to peroxisomes, as well as to mitochondria, lipid droplets, and the endoplasmic reticulum. A comprehensive *Trypanosoma* PEX19 interactome analysis uncovered PEX38 as a novel and essential Euglenozoa-specific peroxin. PEX38 contains distinct domains that bind the co-chaperone Hip and the PEX3-binding motif of PEX19, suggesting a role in stabilizing membrane protein and preventing premature membrane docking. PEX38 illustrates functional repurposing in organelle biogenesis. It originated from a remnant of the GET/TRC pathway, typically responsible for the targeting of tail-anchored proteins to the endoplasmic reticulum. While most components of this machinery are absent in *Euglenozoa*, PEX38 has been retained and adapted to mediate peroxisomal membrane protein targeting. This evolutionary adaptation is unique to *Euglenozoa*. Because the PEX19–PEX38 interaction is essential for parasite viability and PEX38 has no human homologs, this complex is a promising therapeutic target against trypanosomatid parasites.

## Introduction

Approximately 30% of the proteome comprises of membrane proteins, which must be accurately sorted to their target membranes and inserted with the correct topology. This requires elaborate machineries which are further complex in eukaryotes with membrane bound subcellular organization. Membrane proteins are targeted to their destination membranes either co-translationally or post-translationally. Co-translational transport prevents the release of hydrophobic membrane proteins into the cytosol, where they could aggregate or become toxic. However, certain membrane proteins, such as tail-anchored (TA) proteins, require post-translational targeting. To prevent mistargeting or aggregation, some membrane proteins are translated in close proximity to their target organelle, a process known as organelle-associated translation^1^.

PEX19 acts as cytosolic receptor for newly synthesized peroxisomal membrane proteins (PMPs), which are recognized through their membrane peroxisome targeting signal (mPTS). Its flexible N-terminal region contains a PEX3 binding motif that is essential for docking at the peroxisomal membrane, whereas the structured C-terminus mediates binding to the PMPs^2^. PEX19-bound PMPs are targeted to the peroxisomal membrane through binding to the docking factors PEX3 and PEX16^3,4^. Subsequently, the PMP cargo is inserted into the membrane, and PEX19 is released to engage in a new import cycle. Despite this general model, the exact mechanisms governing PMP insertion and receptor release remain poorly understood.

PEX19 has been proposed to function not only as a receptor but also as a chaperone for its PMP cargo, based on two main observations, i) PMPs are unstable in the absence of PEX19, and ii) *in vitro* synthesized PMPs are stabilized in presence of recombinant PEX19^5,6^. However, this chaperone-like role warrants re-evaluation. Notably, PMPs also exhibit instability in the absence of PEX3 and/or PEX16, suggesting that PMP degradation may result from mislocalization to the cytosol in the absence of peroxisomal membranes, rather than from the absence of PEX19 per se^7^. Furthermore, the reported stabilization of the *in vitro* synthesized PMPs in the presence of recombinant PEX19 could also include contributions from the unknown components of reticulocyte lysates used in the study. Moreover, PEX19 lacks sequence or structural similarity to canonical chaperones/HSPs, suggesting that PEX19 may not act as a chaperone in the classical sense. PEX19 client proteins contain single or several transmembrane domains, with TMDs located near N- or C-termini or dispersed throughout the cargo proteins. Chaperones can function as holdase (ATP-independent, preventing accumulation of misfolded proteins), foldase (ATP-dependent, actively transition misfolded proteins back to their native conformation) or disaggregase^8^. It is likely that PEX19 only functions as holdase, preventing the aggregation of membrane proteins, as shown for huntingtin^9^. Active folding of the membrane proteins has not been demonstrated for PEX19. For targeting PMPs with multiple transmembrane domains, PEX19 is unlikely to function alone.

PEX19 primarily targets membrane proteins to peroxisomes, but it also facilitates the targeting of proteins to mitochondria, ER and lipid droplets^5,10^. Mutations in PEX genes lead to peroxisome biogenesis disorders (PBDs), which result in severe neurological abnormalities and early death. These defects are exacerbated by damage to the mitochondrial integrity caused by mislocalization of PMPs to the mitochondria in the absence of peroxisomal targeting^7^.

Furthermore, pathogens, including viruses, can exploit the peroxisomal import machinery to enhance their survival, for example by directing viral protein to peroxisomes via interaction with PEX19^11,12^. Therefore, dissection of the early steps in PEX19-PMP targeting warrant investigation.

All trypanosomatid parasites possess a single membrane-bound peroxisome-related organelle known as glycosome^13,14^. This unique organelle compartmentalises glycolytic enzymes and is involved in essential metabolic pathways, including gluconeogenesis, pentose phosphate, purine salvage, pyrimidine biosynthesis, fatty acid elongation, and sterol biosynthesis^15^. Given the critical role of glycosomes in trypanosomatid parasites, their biogenesis is an important therapeutic target.

In this study, we conducted a comprehensive analysis of the *Trypanosoma* PEX19 interactome by label free quantitative mass spectrometry. Among the top interactors identified were PEX38, a *Euglenozoa* specific peroxin, and the Hsc70 interaction protein (Hip). Using peptide arrays, biochemical assays and high-resolution NMR structural analysis, we demonstrate that PEX38 directly binds to the PEX3 binding motif of PEX19 and serves as a molecular bridge between PEX19 and HiP. A high-resolution NMR structure of the complex reveals molecular details for the interaction further supported by mutational analysis. These findings reveal a previously unrecognized mechanism by which PEX19 may access cytosolic chaperones for stabilizing membrane proteins and facilitating their glycosomal import. Interestingly, most canonical GET pathway components are absent in *Euglenozoa*. PEX38 appears to represent a repurposed remnant of this pathway, remodelled to function as a co-chaperone that bridges the PEX19-PMP complex with the chaperone machinery. We show that both PEX38 and its interaction with PEX19 are essential for parasite viability. Notably, this interaction is absent in the human counterpart making PEX38 a promising candidate for structure-based drug development against trypanosomatid parasites.

## Results

### PEX19 interactome analysis identifies PEX38 as predominant cytosolic binding partner

To identify interaction partners of PEX19, we performed affinity pull-down assays using recombinant GST-tagged *T. brucei* PEX19 (full-length i.e., FL or lacking PEX3 binding region i.e., ΔN30; purification profile shown in **Fig. S1**) in combination with cytosol or solubilized organellar fraction obtained from *Trypanosoma* parasites (**Fig. 1a**; **Fig. S2a**). PEX19 and the bound proteins, eluted using thrombin cleavage, were analysed by SDS-PAGE with Coomassie (**Fig. 1b**) and silver staining (**Fig. S2b**), which revealed potential binding partners as additional bands in comparison to the GST control. Notably, a prominent cytosolic binding partner of ∼38 kDa was detected in the Coomassie stained eluate of PEX19^FL^ but was almost absent in the eluate of PEX19^ΔN30^ (**Fig. 1b**). This protein was unlikely to be PEX3 itself, since it is a membrane protein of a predicted molecular size of 52 kDa.

**Figure 1:**
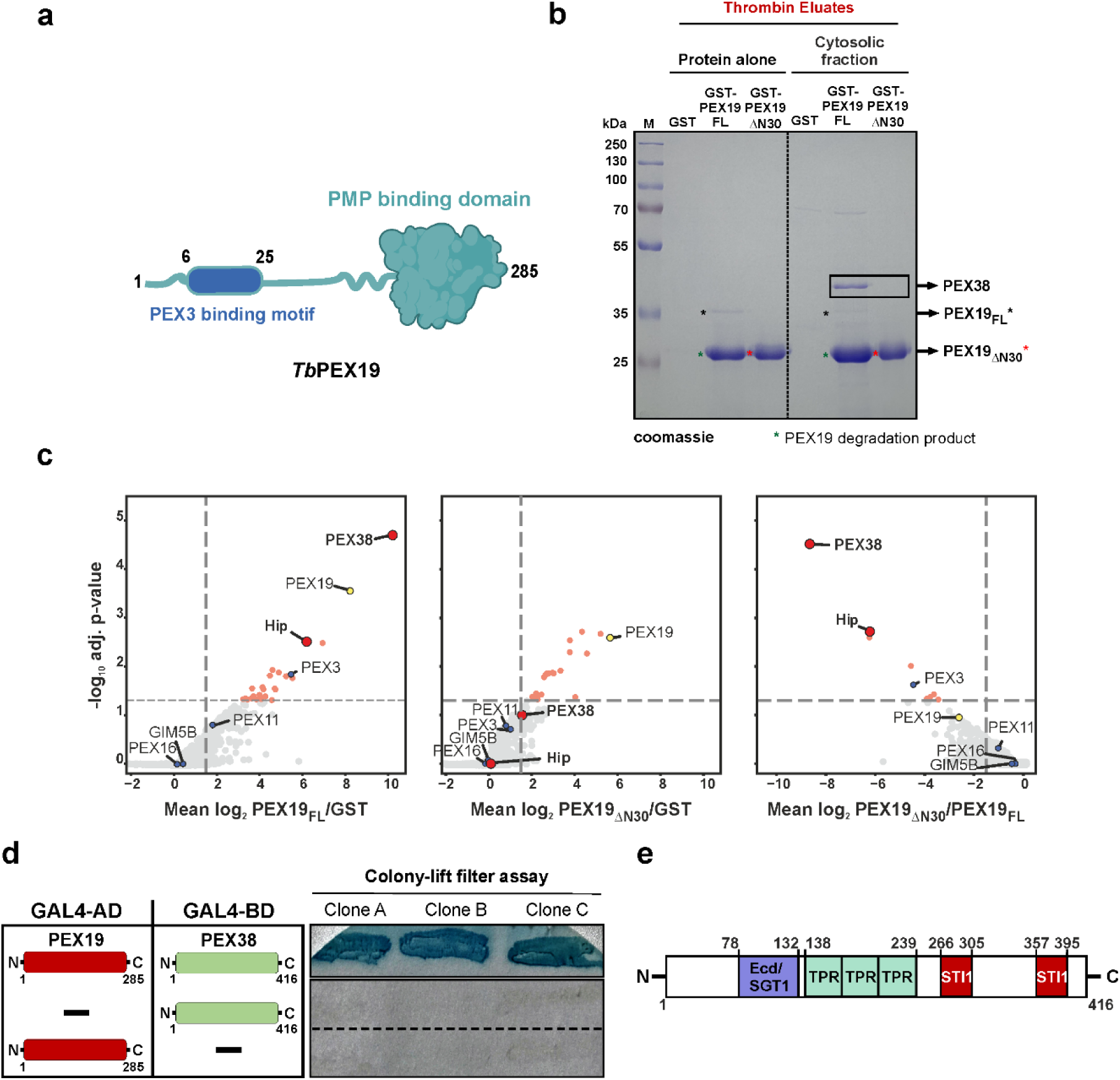
Identification of PEX38 as a dominant binding partner of PEX19. **a)** PEX19 Domain architecture depicting a globular C-terminal PMP binding domain (green) and an unstructured N-terminal domain containing PEX3 binding motif (blue). **b)** Affinity pull-down of *Trypanosoma* cytosolic proteins interacting with *Tb*PEX19. The thrombin eluates were analysed by SDS-PAGE followed by Coomassie staining. PEX38 is highlighted as a dominant putative *Tb*PEX19 binding partner by a black box. **c)** Comparison of the cytosolic interactomes of GST- PEX19FL, GST-PEX19ΔN30, and GST control analysed by Affinity Purification Mass Spectrometry (AP-MS). Ratios (left to right) of PEX19FL/GST, PEX19ΔN30/GST, and PEX19ΔN30/PEX19FL were calculated from quantified iBAQ MS intensities, and a rank sum method was used for the identification of fold differences (x-axis, in log2 scale) in the detected proteins and determination of adjusted p-values (y-axis, in -log10 scale). Highlighted proteins are bait PEX19 (yellow), known glycosomal membrane proteins (blue), PEX38, and HiP (red), proteins with significant fold change (light red). Vertical dashed lines indicate a log2 ratio of greater than 1.5 (left, middle) or less than 1.5 (right) respectively (n ≥ 2 replicates), and the horizontal line indicates a false discovery rate of 5% (n = 3 biological replicates). **d)** Yeast two-hybrid (Y2H) analysis of the PEX19-PEX38 interaction performed in three biological replicates, each with three different clones, by Colony-lift filter assay. **e)** Domain architecture of PEX38. The ECD (Ecdysoneless) domain, also known as SGT1 (Suppressor of the G2 allele of skp1), TPR (Tetratricopeptide Repeat), and STI1 (STress Inducible 1).

The affinity-purified complexes were analysed by label-free quantitative mass spectrometry (MS) to define the cytosolic and organellar PEX19 interactomes. From the 2361 total identified protein groups, we calculated protein abundance ratios between PEX19 and the GST control (PEX19^FL^/GST), as well as the ratios between the two PEX19 variants (PEX19^ΔN30^/GST and PEX19^ΔN30^/ PEX19^FL^). Proteins significantly enriched with both PEX19 variants from cytosolic (**Fig. 1c**) and organellar fractions (**Fig. S2c**) were determined based on rank-sum statistical analysis (n=3, adj. p value < 0.05, mean ratio > 2.25). Of the 24 potential interaction partners of full-length PEX19 in the cytosol, a protein with TriTrypDB ID ‘Tb427_060045000’ was the most significantly enriched (**Fig. 1c**, left panel; **Table S1**). Orthogonal validation by yeast two-hybrid (Y2H) analysis also revealed a clear interaction of this protein with full length PEX19 (**Fig. 1d**). Based on its strong interaction with PEX19 and subsequent characterization, we designated this protein as PEX38. The two other highly enriched proteins for PEX19^FL^ in the cytosol were Hsc70-interacting protein (HiP) and PEX3.

Similarly, among the 45 potential PEX19 interactors identified in the organellar fraction, PEX38 was the most significantly enriched protein after PEX3 (**Fig. S2c**, left panel). Interestingly, this interaction was abolished in the truncated PEX19 variant lacking the N-terminal 30 amino acids (PEX19^ΔN30^) in both cytosol (**Fig. 1c**, middle panel) and organellar fractions (**Fig. S2c**, middle panel), suggesting that PEX19 requires its extreme N-terminal region for binding to PEX38. To further validate this observation, we compared the protein abundance between the two PEX19 variants and found that both PEX38 and HiP were depleted in the cytosolic fraction of PEX19^ΔN30^ relative to full-length PEX19 (PEX19^FL^) (**Fig. 1c**, right panel). Likewise, PEX38 and known PMPs like PEX3 and PEX16 were depleted in the organelle-enriched fraction (**Fig. S2c**, right panel). The top binding partners of both variants in the cytosolic and organellar fractions are listed in **Table S1**.

Bioinformatic analysis of *Tb*PEX38 using BLAST revealed yeast Sgt2 and human SGTA/SGTB as its closest homologs. Both PEX38 and human or yeast SGT proteins share a Tetratricopeptide-like helical superfamily domain and belong to the SGT (Small glutamine-rich tetratricopeptide repeat-containing) protein family (**Fig. 1e**, **Fig. S3a**). However, *Trypanosoma* PEX38 protein displays notable differences compared to the human and yeast SGT proteins. The N-terminal ‘SGTA homodimerization domain’ (InterPro ID: IPR032374, shot name: SGTA_dimer) conserved in SGT2/SGTA/SGTB is not detected in PEX38, and glutamine-rich region at the C-terminus, a characteristic name-giving feature of SGT proteins is also absent in PEX38. Instead, *Tb*PEX38 contains a region with similarity to the Ecd (Ecdysoneless) domain found in human Ecd (hECD), also known as SGT1 (Suppressor of GSR2 1) (**Fig. 1e**). *Tb*PEX38 also contains two STI1 domains, which represent heat shock chaperonin-binding regions, a feature not detected in human *Hs*SGTA/B and yeast *Sc*Sgt2 via InterPro scan. Overall, *Tb*PEX38 shares relatively low sequence identity (∼24-31%) with *Hs*SGTA/B or *Sc*Sgt2 with the highest conservation localized to the TPR domain (**Fig. S3b, c**). Consistent with these structural or domain differences, Y2H analysis showed no detectable interaction between the host *Hs*PEX19 and SGTA, while a weak interaction was observed for *Sc*PEX19 and *Sc*Sgt2 (**Fig. S4a**). Although *Tb*PEX19 shares relatively low sequence homology with host *Hs*PEX19, its PEX3-binding motif is conserved^16^. Therefore, *Tb*PEX19 was also tested with host *Hs*SGTA and *Sc*Sgt2, but no interaction was observed (**Fig. S4b**). These distinctive structural features and experimental results underscore the uniqueness of PEX38 protein to trypanosomatids.

### Characterization of PEX19-PEX38 protein-protein interaction (PPI)

Proteomic and Y2H analysis revealed a clear interaction between full length *Tb*PEX19 and *Tb*PEX38 (**Fig. 1**). To further delineate the PEX19-PEX38 binding interface, we employed synthetic peptide arrays spanning the sequences of each protein. These arrays were probed with recombinantly purified GST-tagged versions of the interacting partner, followed by antibody detection of the probe-bound interaction peptide spots. On the *Tb*PEX19 array, two spots corresponding to the N-terminal region spanning amino acid residues 1-25 and 6-30 of PEX19 indicated potential binding by PEX38 (**Fig. 2a**, middle panel), in contrast to the GST control, which showed no binding in this region (**Fig. S5a**). This N-terminal region (1-25 residues) of PEX19 is also known to mediate the interaction with PEX3 at the peroxisomal/glycosomal membrane, a critical docking step for targeting and membrane insertion of PMPs across different organisms (**Fig. 2a**, scheme on right)^16^. Similarly, using the PEX38 peptide array, we identified four potential PEX19 binding sites (BS1-BS4) in PEX38 (**Fig. 2b**, middle panel), compared to the control GST (**Fig. S5b**). The identified PEX19 binding sites in PEX38 are located in regions comprising amino acid residues 69-93, 89-107, 115-133 and 141-159. Notably, the first three binding sites (BS1-BS3) are located within the Ecd/SGT1 domain, which is only present in PEX38 (**Fig. S3a**), while the fourth binding region (BS4) is situated in the TPR domain (**Fig. 2b**, scheme on right).

**Figure 2:**
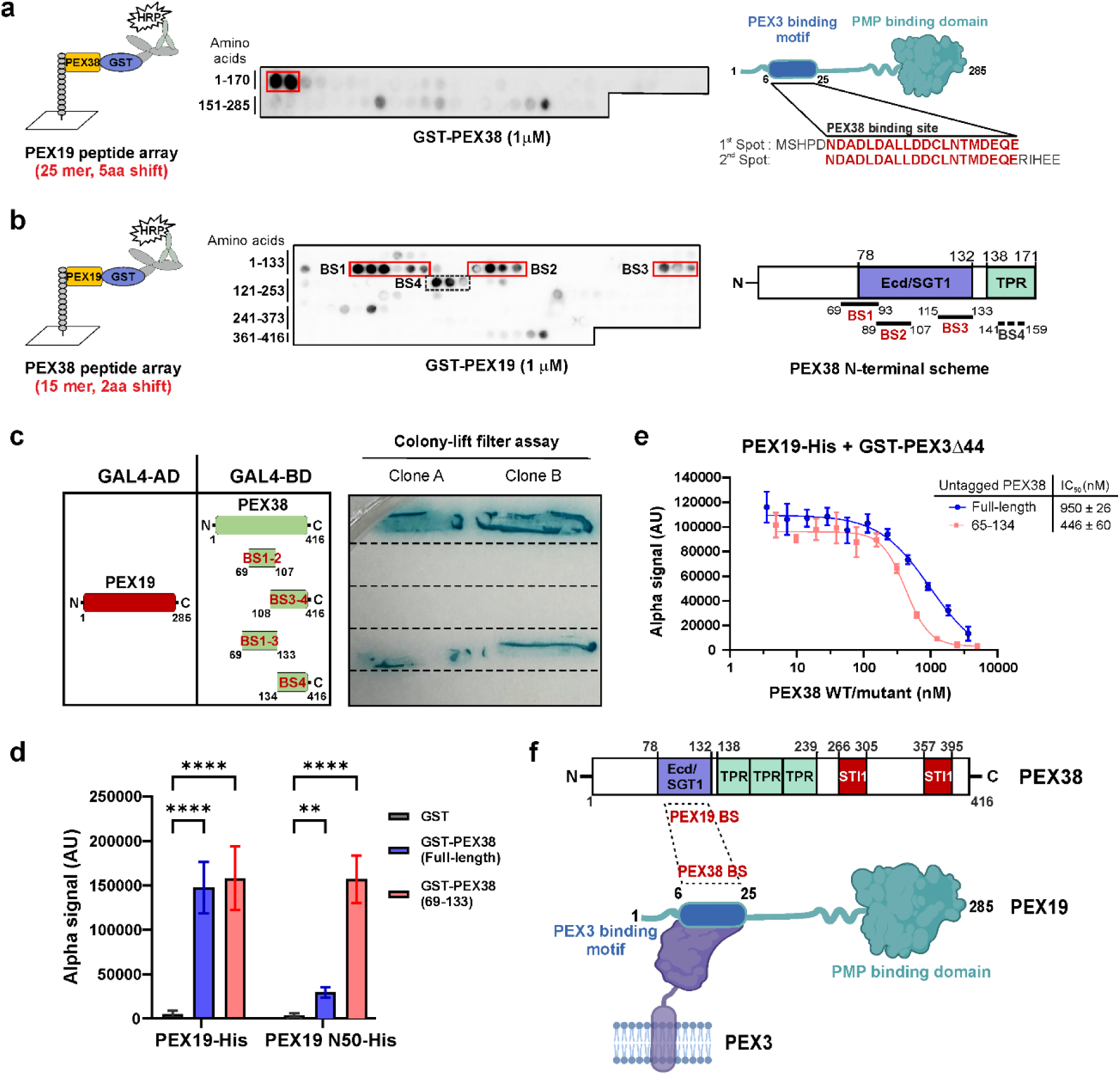
Cytosolic PEX38 and membrane docking protein PEX3 bind to the N-terminal helix of PEX19. **a)** Scheme of PEX19 Peptide array analysis (left). *Tb*PEX19 peptide array was probed with GST (control, **Fig. S5a**) or GST-*Tb*PEX38, and peptide spots bound by the probe were detected by chemiluminescence (middle). Schematic representation of the identified binding site (right), which overlaps with the known PEX3 binding motif. **b)** Scheme of PEX38 peptide array analysis (left). *Tb*PEX38 peptide array was probed with GST (control, **Fig. S5b**) or GST-PEX19, and peptide spots bound by the probe were detected by chemiluminescence (middle). Four *Tb*PEX19 interacting regions identified in PEX38 are shown in red boxes (BS1-BS3) and dotted box (BS4). Schematic representation of the identified PEX19 binding regions in the N-terminal domain of PEX38 (right). **c)** Validation of the PEX19 binding sites in PEX38 using Y2H assay. Colony-lift filter assay with two independent clones is shown (negative controls, **Fig. S5c**). **d)** AlphaScreen-based binding assay for the indicated constructs of recombinant PEX19-His with GST-PEX38 (full-length or 69-133) or GST as a negative control. Statistical analysis was performed using 2-way ANOVA with Šídák’s multiple comparisons test. **** p<0.0001; ** p=0.0012. Error bars represent standard deviations from the mean of n = 3 independent biological replicates, each with six technical replicates. **e)** AlphaScreen-based competition assay to demonstrate that untagged PEX38 (full-length or 65-134) can displace GST-PEX3Δ1-44 from binding to PEX19-His in a dose-dependent manner, with IC50 values of 950 nM and 446 nM, respectively. **f)** Schematic summary of the PEX19-PEX38 interaction mapping and known PEX19-PEX3 interaction.

The PEX19 binding regions identified in PEX38 via peptide array analysis were further tested by Y2H analysis (**Fig. 2c**). The individually tested constructs, PEX38^69-107^ (BS1-BS2), PEX38^108-416^ (BS3-BS4), and PEX38^134-416^ (BS4), did not interact with PEX19 in Y2H. However, a construct PEX38^69-133^ comprising the conserved Ecd/SGT1 domain, which contains BS1-BS3, showed a clear interaction with PEX19, comparable to that observed with the full-length PEX38 (**Fig. 2c**), with negative controls confirming the absence of autoactivation (**Fig. S5c**). BS4 could not be conclusively validated and may require additional experimental strategies to determine its contribution to the interaction.

As an independent approach, we assessed the PEX38-PEX19 interaction using a semi-quantitative AlphaScreen-based assay (**Fig. 2d**). The assay revealed a strong interaction between full-length PEX19 (PEX19^FL)^ and both full-length PEX38 (PEX38^FL^) and the truncated PEX38^69-133^. A short N-terminal fragment of PEX19 (PEX19^1-50^) showed 10 times weaker interaction with PEX38^FL^. However, the same PEX19^1-50^ fragment displayed robust interaction with PEX38^69-133^, which bound PEX19^FL^ and PEX19^1-50^ with comparable affinity. *In vitro* pull-down experiments with same proteins/fragments qualitatively confirmed these interactions (**Fig. S5d**). Taken together, these data identify and validate PEX38-PEX19 protein-protein interaction.

Docking of the PMP-bound PEX19 to the glycosomal membrane relies on the binding between PEX3 and the N-terminal region of PEX19. Since both PEX38 and PEX3 bind to same region in PEX19 (**Fig. 2a**, right panel scheme), we investigated whether these two proteins compete for PEX19-binding. To this end, we performed an AlphaScreen-based displacement assay using equimolar concentrations of PEX19 and PEX3^Δ1-44^ proteins, while titrating increasing amounts of untagged PEX38 (FL or 65-134 constructs). Both PEX38 constructs were able to displace PEX3 from PEX19 in a dose-dependent manner (**Fig. 2e**). Quantitively, ∼950 nM of PEX38^FL^ and ∼446 nM of PEX38^65-134^ were required to displace 30 nM of GST-PEX3Δ44, suggesting that PEX3 has a higher affinity for PEX19 than PEX38. This higher affinity is likely due to additional PEX3-binding regions outside of the PEX19 N-terminus, as previously reported^16–18^.

In conclusion: By combining cell fractionation with pull-down and quantitative MS analysis, we identified a previously unknown cytosolic PEX19-binding protein, PEX38, which interacts with the N-terminal region (1-30) of PEX19, the same region involved in binding of PEX19 to the membrane docking factor PEX3.

### An amphipathic helix from PEX19 interacts with a helical bundle of PEX38

In previous peptide array experiments, we identified the N-terminal amino acid segments 1-25 and 6-30 within PEX19 to interact with a small domain of PEX38^65-134^ (**Fig.2f**). To map this interaction and for high-resolution structural analysis, we generated a PEX19^1-50^ construct with a C-terminal SGGY extension to facilitate detection at 280nm. We performed isothermal titration calorimetry (ITC) and NMR titrations to provide quantitative binding affinities and map the interaction at resolution and finally determined the structure of the PEX38/PEX19 complex.

NMR titration experiments were conducted using ^15^N-labeled PEX19^1-50^ titrated with unlabelled PEX38^65-134^, and reciprocally unlabelled PEX19^1-50^ into ^15^N-labeled PEX38^65-134^. Both setups revealed an intermediate to slow exchange binding regime, as evidenced by a decrease followed by an increase in peak intensities with increasing ligand concentration (**Fig. 3a, b**; **Fig. S6a, b, c**). This behaviour is consistent with the dissociation constant (*K*^D^) of approximately 3 µM determined by ITC (**Fig. 3c**; **Table S2**). Notably, titration of unlabelled PEX19^1-140^ onto ^15^N-labeled PEX38^65-134^ induced similar effects to those observed in the PEX38 spectra, confirming that the binding site is indeed covered by PEX19^1-50^ (**Fig. 3a**; **Fig. S6a, b**). However, residue F73 exhibited significant line broadening during titration with PEX19^1-140^, suggesting a different role in binding to PEX19 full-length (FL) compared with PEX19^1-50^ (**Fig. S6b**).

**Figure 3:**
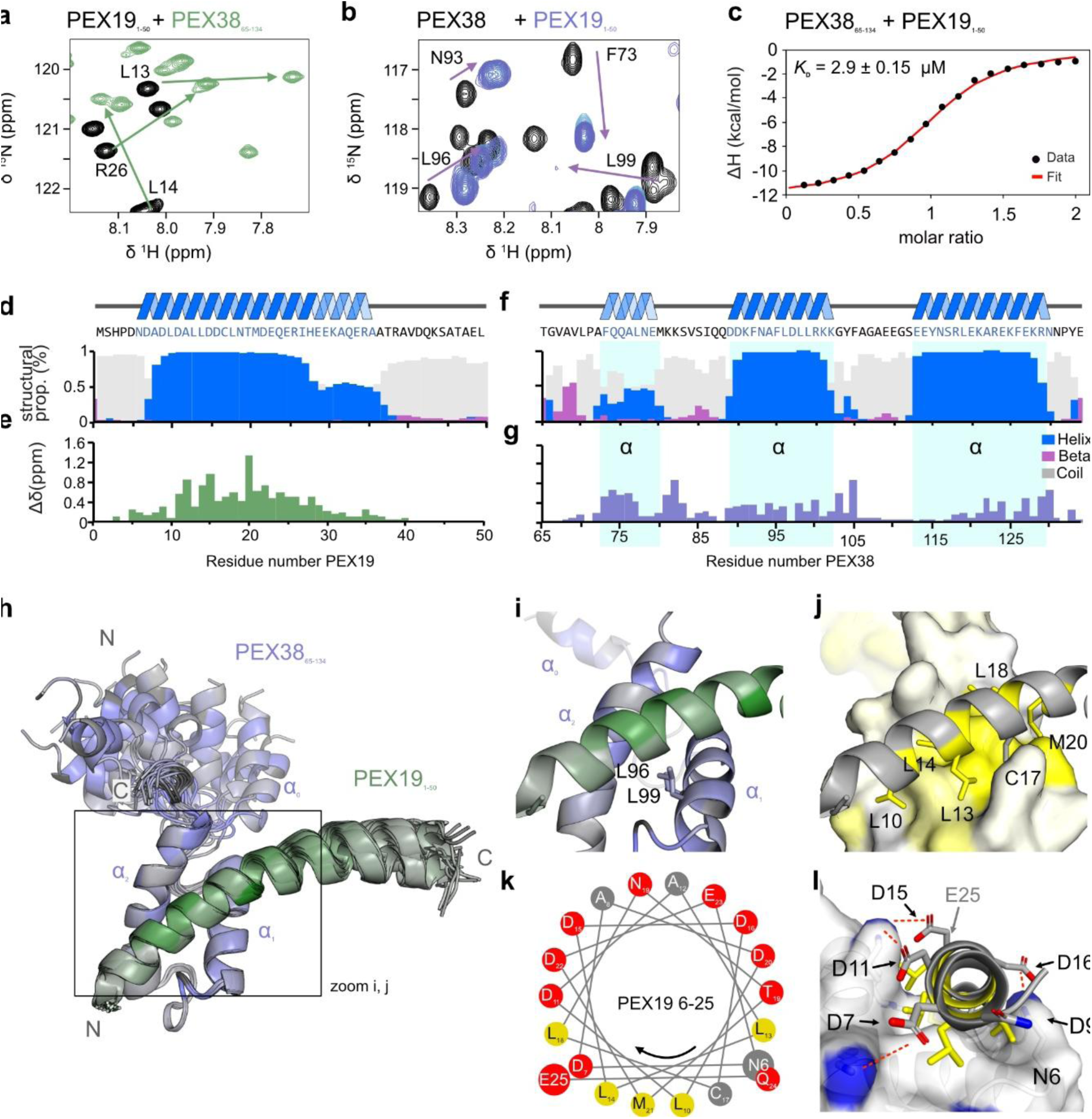
The induced amphipathic helix of PEX191-50 binds to the hydrophobic core of PEX3865-134 helical bundle. **a)** NMR titration of ^15^N labelled PEX191-50 with unlabeled PEX3865-134 and **b**) ^15^N labelled PEX3865-134 with unlabeled PEX191-50 shows large chemical shift perturbations and line-broadening, indicating a strong interaction between the proteins (full spectra shown in **Fig. S6a-c**). **c)** ITC experiments showing titration of PEX38 with PEX19. The experiment was conducted in technical triplicates. **d, f)** Secondary structure propensities of PEX191-50 and PEX3865-134 were predicted using TALOS-N ^41^, based on the secondary chemical shifts (**Fig. S6f, k**). The propensities for α-helix, β-sheet, and coil structures are shown in blue, magenta, and gray, respectively. The helical regions are also visualized above and mapped onto the PEX19 and PEX38 sequences. **e, g)** Chemical shift perturbations (Δδ in ppm) of PEX19 saturated PEX38 or PEX38 saturated PEX19 plotted on the sequences of each protein. **h)** Chemical shift perturbations (from panels **f** and **g**) mapped on an overlay of the 10 best-scored structures with trimmed flexible regions (PEX3865-69 and PEX1939-50). **i)** A zoomed-in view of the first structure highlights the key positions of leucines L96 and L99, which are crucial for the interaction between PEX38 and PEX19. **j)** Surface representation of PEX38 with indicated hydrophobicity ^42^ displayed as white to yellow gradient. Residues from PEX19 L10, L13, L14, C17, L18, M20 facing the hydrophobic surface of PEX38 are represented as colored sticks. **k)** Schematic representation of the PEX19 amphipathic helix 6-25 flanked by N6 and E25. **l)** A front view of the PEX19 amphipathic helix reveals charged residues D7, D11, D15, D16, D22 (E25 is not visible here, please see Fig. S7c) on the hydrophilic side, providing electrostatic interactions with positively charged lysine and arginine residues from PEX38. These contacts are represented by dashed red lines, and the amino groups of the PEX38 lysines are highlighted as blue surfaces. A complete representation is shown in Fig. S7c.

To gain insight into the secondary structure and confirm the binding interface at residue resolution, we recorded triple resonance experiments and assigned the backbone resonances of PEX38^65-134^ and PEX19^1-50^ in both their free and complex states. Secondary chemical shifts (Δδ^13^Cα-Δδ^13^Cβ), obtained from backbone assignments, indicate α-helical and β-strand secondary structure. In the free state, PEX19^1-50^ is largely unstructured, with only regions of low helical propensity (**Fig. S6d, e**). Remarkably, when bound to PEX38^65-134^, PEX19^1-50^ adopts a stable helical structure in the region of amino acid residues 6-27 and a less populated helix from 27-37, while the remaining residues remain unstructured (**Fig. 3d**; **Fig. S6f**). This binding-induced helix formation is further supported by strong chemical shift perturbations of the PEX19^1-50^ amide resonances when compared to free PEX19^1-50^ (**Fig. 3a, e**).

The secondary structure analysis of free PEX38^65-134^ highlights three helical regions α^0^, α^1^ and α^2^ located at amino acid residues 72-79, 90-102 and 113-129, respectively. (**Fig. S6 g, h**). The (α^0^) helix comprises of about two turns, and is about 50% populated, while the longer α^1^ and α^2^ helices, spanning 3 and 5 turns respectively, are fully populated (**Fig. S6 g, h**). Upon binding PEX19^1-50^, the helical propensity of α^1^ and α^2^ increases slightly (**Fig. 3f, g**; **Fig. S6j, k**), while the helical propensity of α^0^ decreases (**Fig. S6 h, k**). To assess backbone flexibility on the pico- to nanosecond timescale, we recorded {^1^H}-^15^N heteronuclear NOE experiments, which showed increased rigidity in helices α^1^ and α^2^ but not in α^0^, upon binding of PEX19^1-50^ (**Fig. S6 i, l**).

Next, we determined the NMR solution structure of the PEX38^65-134^-PEX19^1-50^ complex (**Fig. S7a**), utilizing full side-chain assignments and multiple NOESY experiments. This provided 2752 proton-proton distance restraints, including 224 intermolecular long-range restraints (**Fig. S7b, c; Table S3**). A comprehensive overview of the structure calculation statistics and quality assessment is provided in **Table S3** and **Table S4**, respectively. Analysis of the calculated complex confirmed that the α^0^ region of PEX38^65-134^ and the disordered region of PEX19 (37-50) do not contribute to the complex structure, as shown by transparency or omission in subsequent figures (**Fig. 3h**; **Fig. S7b**). The structures reveals two key leucine residues, L96 and L99, in the hydrophobic core of PEX38^65-134^, which exhibited significant chemical shift perturbations and line-broadening (**Fig. 3a, i**). L96 mediates contacts with the second helix (α^1^) and a loop region, while L99 resides at the hydrophobic binding interface between PEX38 and PEX19, explaining the observed perturbations and line-broadening. The hydrophobic surface of PEX38 (**Fig. 3j**) is occupied by hydrophobic residues L10, L13, L14, C17, L18, and M20 from the amphipathic helix formed by PEX19^6-25^ (**Fig. 3j, k**). On the hydrophilic face of the amphipathic helix, flanked by Asn6 and Glu25, multiple aspartates (D7, D11, D15, D16) and one glutamate (E23) confer a negative charge, enabling additional salt bridge contacts with positively charged side chains of neighboring Lys residues (K102, K121, K125) and Arg129 on the PEX38 side (**Fig. 3k, l**; **Fig. S7c**).

Building on our structural insights, we sought to disrupt the PEX19-PEX38 interaction through targeted mutagenesis. Given the numerous intermolecular NOEs between the α^1^ helix of PEX38^65-134^ and PEX19^1-50^, we performed a ’proline walk’ mutagenesis on a synthetic 15- mer peptide spanning the α^1^ (98-102) (**Fig. S8a**). Notably, substituting any of the central residues FLDLL (95-99) with proline completely abolished its interaction with PEX19 (**Fig. S8b**). This was further confirmed by the Y2H assay, where we introduced proline substitutions to disrupt the helical structure (L96P or LDL95-98PPP), or aspartic acid substitution to disturb the hydrophobic core (L96D). Wild-type PEX38^FL^ as well as PEX38^69-133^ interact with PEX19, while all three mutations of PEX38 completely abolished this interaction (**Fig. S8c, d**). These results corroborate our structural data and underscore the critical role of the helical structure and the hydrophobic core for PEX19 binding (**Fig. 3i, j**).

### Cytosolic PEX38 is required for glycosome biogenesis, and its interaction with PEX19 is essential for parasite survival

PEX38 lacks predicted transmembrane domains or significant hydrophobic regions, suggesting that it is a soluble protein localized either in the cytosol or in the organelle matrix. To determine its subcellular localization, we performed subcellular fractionation of procyclic form *T. brucei* cells. Cells were mechanically ruptured and subjected to differential centrifugation to separate the cytosolic fraction (supernatant) and the organelle-enriched pellet. The pellet was subsequently fractionated by density gradient centrifugation. Immunoblot analysis revealed that PEX38 was almost entirely released in the cytosolic supernatant of the initial differential centrifugation, similar to the cytosolic marker enolase, supporting the conclusion that PEX38 is primarily a cytosolic protein (**Fig. 4a**). A minor portion of PEX38 migrated to density gradient fractions 19 and 20, which contain mainly mitochondrial markers, but also trace amounts of enolase and glycosomal markers. Mature glycosomes migrate in fractions 12-14, while presence of glycosomal markers in fraction 19 may represent nascent glycosomes. In summary, PEX38 is predominantly cytosolic, with a small fraction potentially associating with immature glycosomes or other cellular compartments.

**Figure 4:**
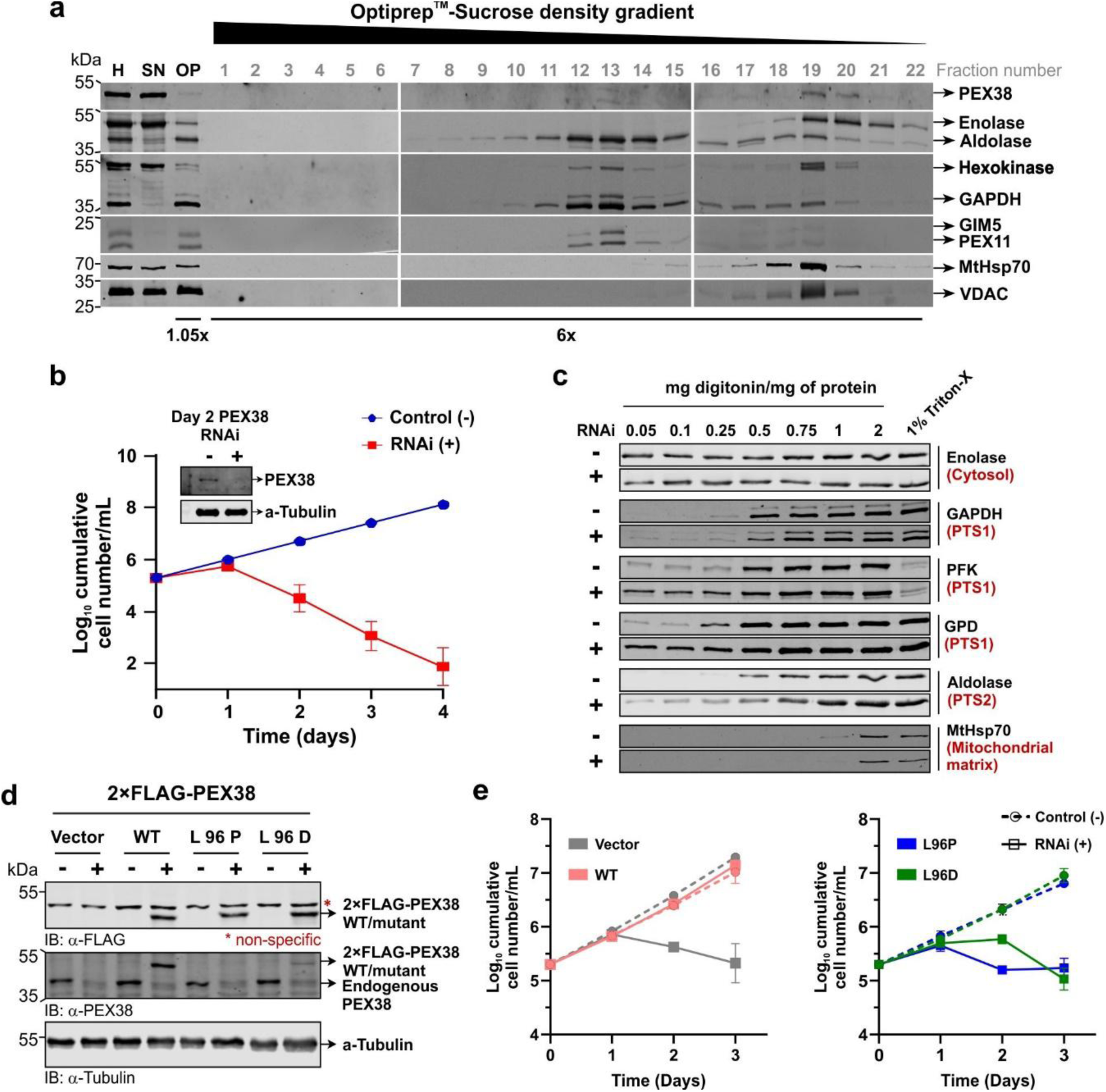
PEX38 is essential for glycosome biogenesis and parasite survival. **a)** Subcellular fractionation of trypanosomes by density gradient centrifugation. Fractions collected from the bottom of the gradient were analysed by immunoblotting, using various subcellular markers. Abbreviations: H, homogenate; SN, supernatant representing cytosolic proteins; OP, organellar pellet. **b)** Cumulative growth curve of the RNAi non-induced control (-, DMSO treated) and RNAi induced (+, Tet treated) PEX38 RNAi cell line. The experiment was performed in triplicate and the error bars represent the mean values with standard deviations. The inset shows an immunoblot analysis with anti-*Tb*PEX38 antibodies to detect the steady state levels of PEX38 in control or RNAi induced trypanosomes. Anti-α-Tubulin serves as the loading control. **c)** Biochemical fractionation of the PEX38 RNAi cell-line. Day 2 PEX38 RNAi non-induced and induced cells were incubated with increasing concentrations of digitonin, centrifuged to yield supernatants, which were analysed immunoblotting with various antibodies. The supernatant of cells incubated with 1% Triton-X 100, a detergent used to dissolve all cellular membranes, is also included. **d)** *in cellulo* functional complementation assay in bloodstream form trypanosomes. The PEX38 RNAi cell line was complemented by ectopic expression of N-terminally FLAG-tagged RNAi-resistant PEX38 derived from a synthetic gene, including the wild-type and mutant forms. Expression of the 2×FLAG-tagged PEX38 wild-type protein and mutants was confirmed by immunoblotting with various antibodies, as indicated. **e)** Cumulative growth curves of the indicated non-induced and RNAi-induced complemented cell lines. PEX38 RNAi cell line transfected with an empty vector (grey), L96P, and L96D (blue and green respectively) exhibited growth defects. In contrast, WT PEX38 (Salmon) restored the growth defect by functional complementation. Error bars represent mean with standard deviations.

We also used *Trypanosoma* cell lines with ectopic, tetracycline-inducible N- or C-terminally GFP-tagged PEX38 to perform immunofluorescence microscopy and assess the localisation of the GFP-tagged PEX38. The expression of these GFP-tagged constructs at correct predicted molecular mass was confirmed by immunoblotting (**Fig. S9a**). Immunofluorescence microscopy was then performed using anti-aldolase antibodies as a glycosomal marker. GFP alone was used as a control to visualize cytosolic distribution. Neither of the overexpressed GFP-tagged PEX38 constructs colocalised with aldolase; instead, the overall cell labelling was like the GFP control, suggesting that the protein is localised in the cytosol (**Fig. S9b**). However, while the overall cell labelling indicates cytosolic localisation, the organelles are not excluded from the labelling, so an additional organellar localisation cannot be distinguished by fluorescence microscopy.

The subcellular distribution of GFP tagged PEX38 was further examined biochemically by cellular fractionation. *T. brucei* cells expressing either N- or C-terminally GFP-tagged PEX38 were treated with a low concentration of digitonin, selectively permeabilizing the plasma membrane and releasing cytosolic contents into the supernatant. Enolase was used as a cytosolic marker, while aldolase served as a marker for glycosomes in the organelle-containing pellet. For both N- and C-terminally GFP-tagged PEX38, the majority of the proteins was detected in the cytosolic supernatant, in agreement with the microscopy data (**Fig. S9c**). Notably, a small fraction of GFP-PEX38 was also detected in the organellar pellet (**Fig. S9c**), suggesting a potential weak association with organelles, similar to what was observed for endogenous PEX38. However, the apparent higher amount of GFP-tagged PEX38 in pellet fraction may also result from overexpression.

### PEX38 is essential for survival of bloodstream and procyclic *T. brucei* forms

To investigate the essentiality of PEX38 for *T. brucei*, we performed a tetracycline inducible RNA interference (RNAi) knockdown of PEX38 gene expression. Cumulative growth of RNAi induced or DMSO treated (negative control) bloodstream-form as well as procyclic-form parasites was monitored for 4 and 9 days, respectively. A severe growth defect is observed for both bloodstream form as well as procyclic form trypanosomes (**Fig. 4b**, **Fig. S9e**). Cell death was evident in the microscopic observation of parasite cultures. The depletion of PEX38 protein level was confirmed by immunoblot analysis (**Fig. 4b**, insets). The results demonstrate that PEX38 is critical for the parasite growth and viability of both bloodstream and procyclic form trypanosomes.

To investigate whether PEX38 depletion affects glycosome biogenesis, we performed immunofluorescence microscopy on *T. brucei* PEX38 RNAi cells (**Fig. S10**). DMSO treated control cell lines showed a characteristic punctate pattern for the glycosomal marker enzymes GAPDH (PTS1 protein) and aldolase (PTS2 protein), consistent with proper glycosome localization. In contrast, RNAi-induced cells exhibited an abnormal glycosomes morphology and a partial mislocalization of both glycosomal matrix markers to the cytosol. However, due to the persistence of pre-existing glycosomes, which stain brightly, the extent of mislocalization was difficult to assess solely by microscopy. To quantitively evaluate this mislocalization, we conducted biochemical fractionation using digitonin permeabilization (**Fig. 4c**). Control or RNAi-induced cells treated with increasing concentration of digitonin were fractionated into cytosolic and pellet fractions. At low concentrations, digitonin permeabilizes the plasma membrane to release cytosol into the supernatant, as evidenced by the release of cytosolic marker enolase. Only at higher concentrations such as 0.5 mg digitonin/mg protein, organelles are permeabilized to release organellar matrix proteins. The analysis revealed that upon PEX38 RNAi, various PTS1- as well as PTS2-containing glycosomal enzymes are released into the cytosol at significantly lower concentrations of digitonin compared to the control. The mitochondrial matrix protein MtHsp70 was not affected by PEX38 RNAi. These findings demonstrate that PEX38 is required for proper glycosome biogenesis and matrix protein import in *T. brucei*.

Like PEX38, partial knockdown of PEX19 also results in a growth defect, demonstrating its essential role^19^. To determine whether the parasite-specific PEX19-PEX38 protein-protein interaction (PPI) identified in this study is essential for parasite survival and thus a potential druggable target, we performed an *in cellulo* functional complementation analysis^20^ in bloodstream form *T. brucei* parasites (**Fig. S11a**). We engineered a PEX38 RNAi cell line to express an ectopic, RNAi-resistant codon-exchanged PEX38 gene under the Tet-inducible promoter (**Fig. S11a, b**). Tet-induction of this double-transfected cell line results in depletion of the endogenous PEX38 and simultaneous expression of the ectopic RNAi-resistant PEX38, which was Flag-tagged to distinguish it from endogenous PEX38. Two controls were used in this assay: a negative control transfected with an empty vector, and a positive control transfected with non-mutated PEX38. Guided by our structural characterisation, we chose the conserved L96 residue within in the Ecd/SGT1 domain of PEX38, which mediates contact with PEX19, for site-directed mutagenesis. We generated two mutants: L96P (disrupting helix formation) and L96D (disrupting hydrophobic interactions) and tested them in the same complementation assay. Prior to growth analysis, immunoblotting analysis confirmed the expression of all ectopic constructs using an anti-Flag antibody (**Fig. 4d**, upper panel) and efficient knock-down of endogenous PEX38 with anti-PEX38 antibodies (**Fig. 4d**, middle panel).The mutant constructs (L96P/D) were clearly detected with Flag-antibodies, but either faintly or not detected by the PEX38 antibody, suggesting these mutations may induce structural changes that compromise antibody recognition. Upon tetracycline induction, the negative control displayed growth defect from PEX38 RNAi, while expression of wild-type PEX38 fully rescued the phenotype, restoring near-normal growth (**Fig. 4e**, left panel). In contrast, neither the L96P nor L96D mutants were able to rescue the parasite growth (**Fig. 4e**, right panel). Taken together, these results demonstrate that the PEX38-PEX19 PPI is essential for *Trypanosoma* parasite survival PEX38 orthologs could be identified in all kinetoplastids as well as in *Diplonema*, which together belong to phylum *Euglenozoa*. As PEX38 appears to be specific to *Euglenozoa*, we extended our study to other clinically relevant kinetoplastids. Multiple sequence alignment of PEX38 orthologs from various *Trypanosoma* and *Leishmania* species and *Diplonema* reveals that the PEX19 binding region in PEX38 is highly conserved (**Fig. S12a**). Notably, the residues L96 and L99, which mediate contacts with PEX19, are conserved across species (**Fig. S12a**). Overall, these sequences share at least 55% identity and more than 63% similarity with *Tb*PEX38 (**Fig. S12b**). To test whether this interaction is functionally conserved, we performed Y2H assays using *L. donovani* PEX19 (*Ld*PEX19) and PEX38 (*Ld*PEX38). As expected, *Ld*PEX19 interacted with wild-type *Ld*PEX38. Importantly, proline substitutions corresponding to the L96 and L99 residues of *Tb*PEX38 abolished the interaction with *Ld*PEX19 (**Fig. S12c**), emphasising the importance of these residues in the clinically relevant organism *L. donovani*. Taken together, these findings demonstrate that the PEX19-PEX38 protein-protein interaction is conserved across trypanosomatid species. Given its functional importance and parasite specificity, the PEX19-PEX38 interaction represents a promising drug target for the development of new therapeutics against trypanosomatid infections.

### PEX38 bridges the PMP import receptor PEX19 with the protein folding machinery via Hip

Previous studies suggested that PEX19 acts as a cytosolic chaperone for newly synthesised PMPs^5^. Overexpressed PEX19 has been demonstrated to stabilise PMPs, while its absence leads to PMP aggregation^5,21–23^. Despite this, the mechanism by which PEX19 mediates PMP chaperoning remains unclear. Along this line, the *Leishmania* ortholog of PEX38, originally termed *Ld*SGT, has been characterized as a co-chaperone that forms a stable complex with the Hsc70-interacting protein (Hip) and associates with Hsp70 and Hsp90 chaperones^24^. To investigate this further, we examined the *Tb*PEX19 interactome in detail and identified a putative Hip homolog as one of the top interactors. Particularly, like PEX38, the putative Hip protein was absent in affinity pull-downs from cytosolic fraction with PEX19^ΔN30^ as bait, indicating that its interaction is dependent on the same PEX19 region (see **Fig. 1b**). This protein retrieves the human Hsc70 interacting protein as the top hit in BLAST search. Henceforth, we will refer to it as the *Trypanosoma* Hip (*Tb*Hip). Domain analysis revealed that *Tb*Hip shares key features with the human Hip, including conserved Tetratricopeptide-like helical and STI1/HOP domains (**Fig. 5a**). However, unlike its human counterpart, *Tb*Hip lacks the N-terminal dimerization domain (Hip_N), suggesting potential differences in complex assembly or regulation.

**Figure 5:**
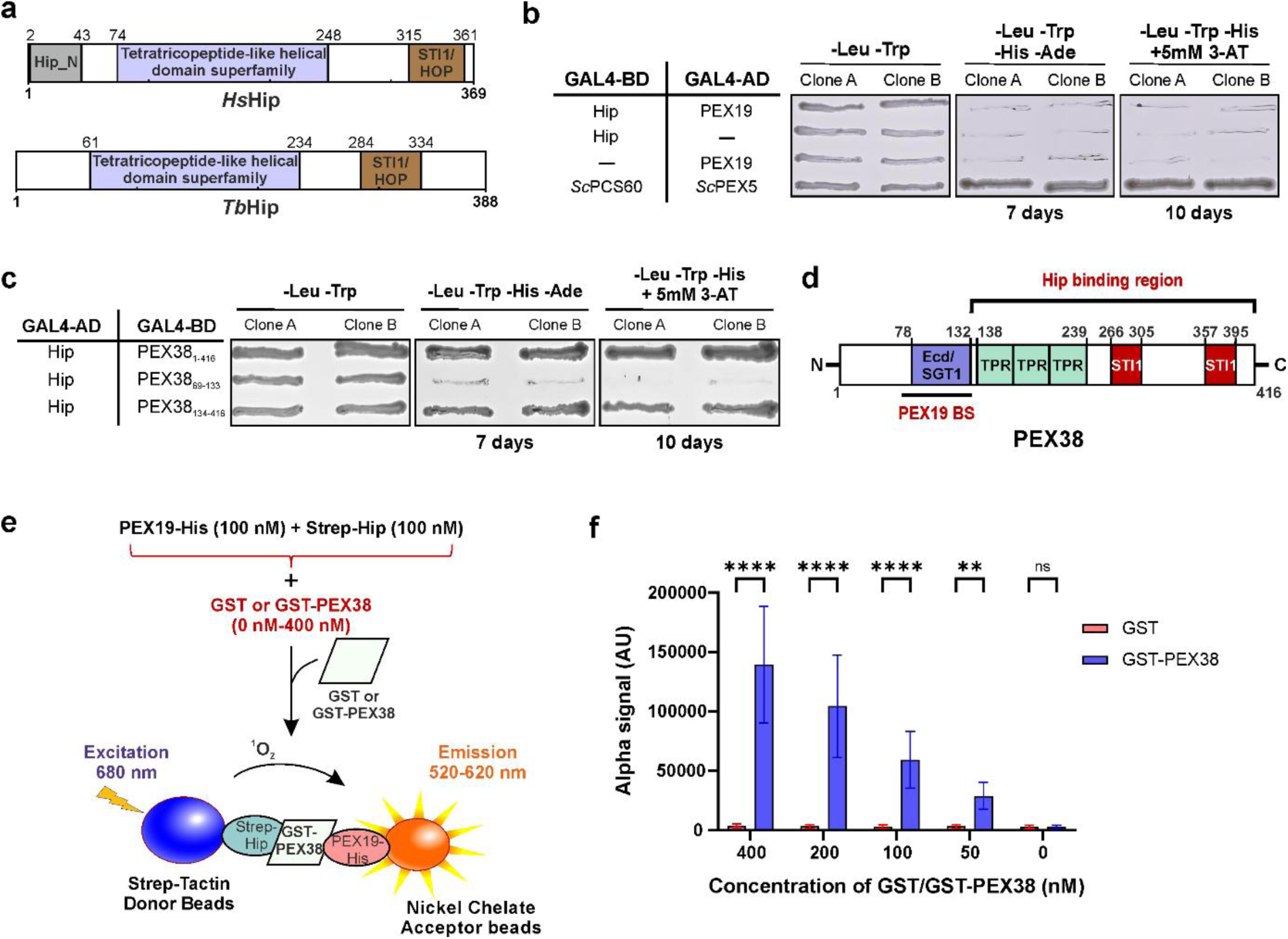
PEX38 connects PEX19 to the chaperone machinery via Hip. **a)** Domain architecture of *Hs*Hip (Hsc70-interacting protein) and *Tb*Hip. **b)** His-auxotrophy-based Y2H assay was performed in the PJ69-4A yeast strain to test the interaction between PEX19 and Hip protein, the second most prominent hit in the PEX19 interactome (see Figure 1b). Two independent yeast clones streaked on -Leu -Trp plates (growth control), -Leu - Trp -His -Ade plates (grown for 7 days) and -Leu -Trp -His + 5 mM 3-amino-1,2,4-triazole plates (grown for 10 days) are shown. Full-length PEX19 and Hip constructs were fused to GAL4 activation (AD) or binding domains (BD), and the interaction was analyzed using a growth-based assay. The *Sc*PEX5-*Sc*PCS60 interaction served as a positive control, while the negative control showed no autoactivation. **c)** Y2H analysis of Hip-PEX38 (FL and variant) interaction. The assay was performed as described in panel **b**, using growth-based readouts to assess interaction. **d)** Schematic summary of PEX19 and Hip interactions with PEX38, showing the distinct binding regions as identified and illustrated in Figures 2b and **5c**. **e)** Schematic representation of the AlphaScreen-based binding assay, which demonstrates the bridging function of PEX38 that results in the ternary complex of PEX19, PEX38, and Hip. **f)** The AlphaScreen assay was conducted using 100 nM of PEX19-His and Strep-Hip, with varying concentrations of GST-PEX38 or GST as indicated. Statistical analysis was performed using 2-way ANOVA with Šídák’s multiple comparisons test. The results showed **** p<0.0001; ** p=0.0013; ns, not significant. Mean values with error bars representing standard deviations from three independent biological replicates are shown, each with six technical replicates.

To further validate the proteomic identification of Hip as PEX19 interactor, we investigated the interaction between PEX19 and Hip using Y2H assays. Unlike PEX38, *Tb*Hip did not interact with PEX19 in the Y2H assay (**Fig. 5b**), suggesting that the association observed in the PEX19 interactome may be indirect. We next tested whether *Tb*Hip interacts with PEX38. All proteins/truncations tested here were verified that they do not show autoactivation (**Fig. S13a**). Full-length PEX38, as well as truncation constructs spanning residues 134-416 that completely lack PEX19 binding site, shows clear interaction with *Tb*Hip. In contrast, PEX38 variants encompassing the PEX19 binding regions (residues 69-133 comprising BS1-BS3) did not interact with Hip (**Fig. 5c**). These findings suggest that Hip interacts with the C-terminal region of PEX38 (residues 134-416), independent of the PEX19- binding region (**Fig. 5d**). Thus, the detection of Hip in the PEX19 interactome is likely mediated through its interaction with PEX38, rather than via a direct association with PEX19. The C-terminal region of PEX38 (residues 134-416), which mediates interaction with Hip, contains two STI1 domains and a region distinct from the PEX19 binding region comprising the Ecd/SGT1 domain. This domain separation may suggest that PEX38 can simultaneously bind to both PEX19 and Hip via non-overlapping regions.

To assess whether disruption of the PEX19-binding site affects PEX38’s ability to bind Hip, we tested PEX38 mutations, which disrupt to PEX19 binding by a Y2H assay. Both wild-type as well as the mutants (mutations L96 to P or D and LDL 96-98 to PPP), showed similar levels of interaction with Hip (**Fig. S13b**). These findings indicate that mutations impairing the binding of PEX38 to PEX19 do not interfere with Hip binding, further supporting the notion that PEX38 has separable binding interfaces for these two partners.

To directly test whether PEX38 can bridge PEX19 and Hip into a ternary complex, we performed an AlphaScreen-based interaction assay using recombinant proteins: PEX19-His, GST-PEX38, and Strep-tagged Hip, with GST also as a control. PEX19-His was pre-incubated with increasing amounts of GST or GST-PEX38, followed by the addition of Strep-Hip. A robust increase in the AlphaScreen signal was observed with increasing amounts of GST- PEX38, indicating the formation of a ternary complex, involving PEX19, PEX38 and Hip (**Fig. 5e, f**). In contrast, no significant signal was detected between PEX19 and Hip in absence of PEX38 or in the presence of GST alone, consistent with the Y2H study (**Fig. 5b**). These data strongly suggests that PEX38 bridges the interaction of PEX19 and Hip to form a ternary complex. This complex may functionally link the targeting of peroxisomal membrane proteins by PEX19 with their folding of stabilization via the chaperon-associated cofactor Hip, highlighting a coordinated mechanism for glycosomal membrane protein biogenesis.

## Discussion

The mechanism by which PEX19 chaperones and targets peroxisomal membrane proteins (PMPs), containing single or multiple TMDs and inserts them in their correct topology into the glycosomal membrane is poorly understood. An earlier study reported that the C-terminal domain of PEX19, although capable of binding PMPs, is insufficient for maintaining the solubility of *in vitro* synthesised PMPs^23^. Moreover, combining separately purified N- and C-terminal domains of PEX19 also failed to stabilize these proteins^23^. A recent study showed that chaperoning of a mammalian TA protein PEX26 by PEX19 involves the amphipathic helix in the C-terminal domain of PEX19, while insertion requires another amphipathic helix in the N- terminal domain^25^. These observations suggested that PEX19 N-terminal domain has additional functions apart from binding to PEX3. Here, we report on the identification and characterization of PEX38, a kinetoplastids-specific peroxin as a key cytosolic bridging factor that links the PMP import receptor PEX19 to the cytosolic chaperone machinery via Hip, a co-chaperone.

In most eukaryotes, PEX19 contains a C-terminal CaaX motif that undergoes farnesylation, enhancing its affinity for PMPs^26^. However, euglenozoan PEX19 proteins, including those in *Trypanosoma* and *Leishmania*, lack the CaaX motif and are not farnesylated. Interestingly, the *Leishmania* ortholog of *Tb*PEX38 (previously termed *Ld*SGT) was shown to act as a co-chaperone, forming stable complexes with the chaperone machinery, including Hsp70, Hsp90, and the co-chaperone Hsc70 interacting protein (Hip)^24^. In our study, proteomic analysis identified Hip and Hsp70 along with PEX38 in the *Trypanosoma* PEX19 interactome, which were significantly reduced in the interactome upon deletion of the PEX38 binding motif, suggesting that PEX38 serves as a mediator for chaperone recruitment to PEX19. This mechanism may compensate for the absence of farnesylation in *Trypanosoma* PEX19 or represent an alternative strategy to engage chaperones during PMP import, since farnesylation is known to allosterically enhance PEX19’s recognition of PMPs by reshaping its binding surface^26^. A similar mechanism has been reported to occur in mitochondria, where receptors TOM22 or TOM70 selectively recognize cytosolic Hsp70 co-chaperones that bind hydrophobic precursor proteins and assist in their transfer from the cytosol to the mitochondrial receptors^27^.

The N-terminal region of PEX38 contains a conserved domain homologous to the Ecd/hSGT1 domain, previously recognized for its role in cell cycle progression in mammals^28^. We demonstrate that this region specifically mediates binding to the N-terminal PEX3-binding motif of PEX19 in both *Trypanosoma* as well as *Leishmania*.

PEX38 shares partial structural similarity with SGT2/SGTA family of co-chaperones from the Guided Entry of TA proteins (GET in yeast)/transmembrane recognition complex (TRC in humans) pathway, a conserved mechanism of targeting TA proteins to the ER^29^, as recognized earlier^24^. SGT proteins, such as Sgt2 in yeast and SGTA in humans capture newly synthesized tail-anchored proteins in the cytosol and target them to the ER membrane, acting in conjunction with other GET proteins or their homologs (reviewed in ^30,31^). However, PEX38 lacks key SGT features, including the conserved SGTA N-terminal homodimerization domain and the C-terminal glutamine-rich region. This, along with its unique domain architecture and demonstrated role in glycosome biogenesis, supports PEX38’s classification as a peroxin rather than a canonical SGT protein. The GET/TRC pathway has been studied mainly in fungi and metazoans, and it is also present in plants and lower eukaryotes, including the human pathogens Plasmodium and Giardia^32^. Interestingly, no clear homologs of canonical GET/TRC pathway components, including SGT2/SGTA, were identified in *T. brucei*. A putative Get1 homolog was detected in *Bodo saltans* (**Fig. S14a**), and a Get3 homolog is present in *B. saltans, T. cruzi* and several *Leishmania* species (**Fig. S14b, c**), although certain species such as *L. tarentolae* encode only fragmented versions. Our bioinformatic analysis (**Fig. 6a**) suggests that while the GET/TRC pathway was likely present in the common ancestor of kinetoplastids, it has been partially or fully lost in several lineages. In this context, PEX38 may represent a lineage-specific adaptation in *Euglenozoa*, functionally replacing components of the GET/TRC-system to facilitate chaperoning of peroxisomal membrane proteins in the absence of canonical machinery.

**Figure 6:**
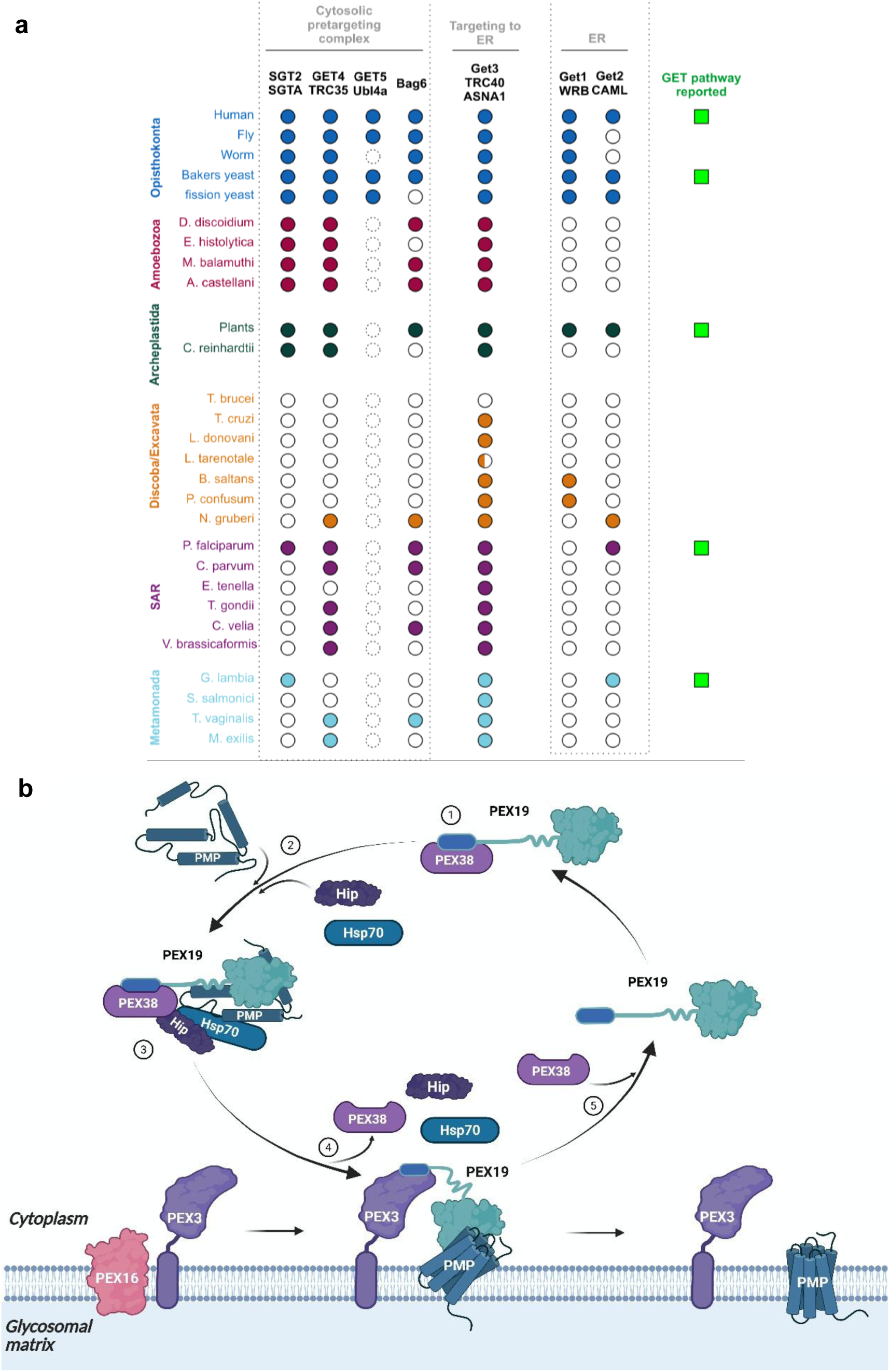
a) Post-translational membrane protein targeting pathway to glycosomes/peroxisomes across organisms. Coulson plot depicting the presence or absence of orthologs of proteins involved GET/TRC pathway. Yeast and human proteins were used to identify orthologs in other organisms. Dotted circles indicate presence of several candidates which cannot be assigned to Get5/Ubl4a unambiguously. Half coloured circle indicates only partial fragment is identified. **b) Proposed model of glycosomal membrane protein import**. Membrane protein import is achieved by a process involving cytosolic receptor, i.e., PEX19, which recognizes the mPTS signal of glycosomal membrane proteins (PMPs). Five distinct steps are conceptualized: **(1) Receptor priming**: PEX38 binds to PEX19, preventing docking of PEX19 at glycosomal membrane without cargo. **(2) Chaperoning of newly synthesized PMPs:** PMPs containing mPTS signals are initially in an unfolded form, bound by chaperones (Hip, Hsp70) preventing their aggregation. **(3) PMP pre-targeting complex formation:** Cytosolic receptor PEX19 with bound PEX38 recognizes the mPTS signal in PMPs which are associated with chaperones. The PEX38-bound PEX19 with folded PMP is ready for docking. **(4) Docking and insertion:** The receptor-cargo complex (PEX19- PMP) docks to the docking protein (PEX3) at the glycosomal membrane. During this step, PEX38 is displaced by PEX3 due to its higher affinity for PEX19 (PMP-complex handover), allowing the insertion of the PMP into the membrane. Displacement of PEX38 accompanies dissociation of Hip and chaperones. Chaperones may assist in active unfolding and folding of PMPs during their insertion in native topology. **(5) Recycling of PEX19:** PEX19 is released back to the cytosol, where it binds to PEX38 and is prepared for the next import cycle of newly synthesized PMP.

Depletion of PEX38 resulted in glycosome biogenesis defects and parasite death, confirming its essential role as peroxin^16,33^. Structurally, our study revealed that the binding region of PEX19 for PEX38 overlaps its PEX3 binding motif, which is crucial for docking PEX19-PMP complexes at the glycosomal membrane^16,18^. Accordingly, our displacement assay revealed a competition between PEX3 and PEX38 for binding to PEX19, but PEX3 binds this region with ∼30-fold higher affinity than PEX38 *in vitro*. However, *in vivo* conditions favour dynamic competition: PEX38 is more abundant in the cytosol than membrane-anchored PEX3 and therefore more likely associated with cytosolic PEX19. PEX3 or PEX38 binding to PEX19 would by dynamic *in vivo* depending on the stage of PMP import cycle and the competition might even serve as functional cooperation. Thus, the interplay between PEX3 and PEX38 may represent a regulated hand-off mechanism during different stages of PMP import, akin to competitive/cooperative interactions seen in other trafficking pathways (e.g., Inp1 vs. PEX19 for PEX3^34^.

The PEX3 binding motif of PEX19 is intrinsically disordered in the absence of PEX3 and is assumed to adopt an α-helical conformation when bound to PEX3^10^. Our NMR analysis demonstrates that PEX38 also induces disorder-to-helix transition in PEX19. Proteins with intrinsically disordered regions play key roles in protein networks, enabling communication through protein-protein interactions, in fact large disordered regions play key roles in peroxisome biogenesis and have been identified in PEX19 homologs but are also found in the N-terminal region of the cargo receptor PEX5 and in the docking protein PEX13^35–37^. Many regulatory proteins have disordered motifs that fold when binding cellular targets, though the underlying mechanisms are unclear^38^.

Based on our findings, we propose a working model for the glycosomal membrane protein import in which PEX38 bound PEX19 interacts with newly synthesized PMPs which are stabilized by Hip and associated chaperones to ensure the proper folding and delivery to the glycosomal membrane (**Fig. 6b**). At the peroxisomal membrane, competition of PEX3 for PEX19 binding will result in the handover of the PMP-loaded PEX19 to PEX3 and in the release of PEX38 together with the associated chaperone machinery. Experimental validation of this model will be critical for understanding the full mechanistic cycle of glycosomal PMP import. Nonetheless, parallels can be seen with recently identified PEX39, which in first step binds with its KPWE motif to PTS2 receptor PEX7, followed by cargo protein and co-receptor binding, and subsequent docking to peroxisomal membrane by binding of KPWE motif of PEX13 and displacement of PEX39 (handover mechanism)^39^. Another study implicates KPWE motif of PEX39 in the last step of PTS2 protein import, where it enables extraction of PEX7 from PEX13 YG phase and prepare it for another round of import cycle^40^.

Importantly, glycosomes are a therapeutic vulnerability (designated as Achile’s heel) of trypanosomatid parasites and represent a validated drug target class for the development of new therapies for devastating trypanosomiasis and leishmaniasis. PEX38 is essential, kinetoplastid-specific, with no equivalent human counterpart. Using *in vivo* functional complementation, we further validated that disrupting the PEX38-PEX19 interaction is lethal to the parasites. Combined with our structural insights, these finding nominate the PEX38-PEX19 binding interface as a promising molecular target for structure-based drug discovery against trypanosomiasis and leishmaniasis.

## Author Contributions

Conceptualization, CK., VK., and RE.; Methodology, CK., SG., HD., and MJ.; Investigation, CK., SG., and HD.; Writing – Original Draft, CK., VK., and RE; Writing – Review & Editing, CK., SG., HD., SO., BW., MS., VK., and RE.; Funding Acquisition, BW., and RE.; Supervision, SO, BW, MS., VK, and RE.

## Acknowledgment

This project work has received funding from the European Union’s Horizon 2020 research and innovation program under the Marie Skłodowska-Curie grant agreement no. 812968 (C.K.K., H.D., B.W., and R.E.). The work was supported by the Deutsche Forschungsgemeinschaft (DFG), grant ER178/17-1 and by Ruhr University Bochum through InnovationsFoRUM (Host Microbe Interactions: IF-009N-22 and IF-018N-22) to R.E, by DFG FOR1905, project number 219314758, SA823/11) to M.S. and R.E. We thank Paul Michels, James D. Bangs, Frank Voncken, and André Schneider for kindly providing various antibodies and the Proteomics Identifications Database (PRIDE) team for data deposition to the ProteomeXchange Consortium. We further thank Frédéric Allain for providing the infrastructure for the final structure calculations. The graphical import scheme was created using BioRender.com.

## Methods

### Cloning

The expression plasmid constructs and cloning strategies for *Escherichia coli*, yeast, and *Trypanosoma* are detailed in **Table S5**, while the oligonucleotide sequences are provided in **Table S6**. The Strep-Hip construct was cloned using the FastCloning method, as described previously^43^. Overlap extension PCR was used to generate the point mutations and gene fragment deletions (PEX38 134-416). Automated Sanger sequencing was employed to verify the sequences of all constructs, mutations, and gene fragment deletions.

For structural characterization, cloning of PEX38 (65-134) and PEX19 (1-50-SGGY) into a pET SUMO expression vector was performed using site-directed ligase-independent mutagenesis (SLIM). To this end, an extended version was applied to implement inserts using the same fashion of short and tail primers. The vector backbone and inserts were amplified by polymerase chain reaction (PCR) amplification using the according short and tail primers (**Table S6**) to generate overlaps with sticky ends. The backbone amplificants and inserts were mixed with a 5-fold molar excess of insert and annealed during the SLIM cycle. The annealed vector was directly transformed into DH10b cells for DNA amplification.

For PEX38 RNA interference, a stem-loop construct was generated using two fragments of the PEX38 gene’s RNAi target region. Fragment 2 (HindIII/MfeI) contained an additional 50-60 base pairs compared to Fragment 1 (SpeI/MfeI), resulting in a step-loop structure. Fragments 1 and 2 were amplified by PCR using primers designed for the RNAi target region using a web-based tool^44^, as specified in **Table S5**. Subsequently, both fragments were subjected to digestion with the common enzyme MfeI, followed by ligation with 250 ng of each fragment. After reaction cleanup, the ligated fragment was further digested with SpeI and HindIII and then cloned into the p2T7-177 vector digested with SpeI-HindIII. For the *in cellulo* complementation assay, a codon-exchanged, RNAi-resistant PEX38 gene with an N-terminal 2×-FLAG tag was custom-synthesized and obtained from GeneCust, France.

### Protein expression and purification

*E. coli* strain TOP10 was used for all plasmid amplifications, with liquid cultures grown at 37°C under continuous shaking in LB medium containing either 100 μg/mL Ampicillin or 50 μg/mL kanamycin. The BL21 *E. coli* strain was used for heterologous expression of recombinant GST, GST-PEX19 full-length (PEX19^FL^) or GST-PEX19 lacking N-terminal 30 amino acids (PEX19^ΔN30^), GST-PEX38 (full-length or 69-133), GST-PEX3Δ44, PEX19-His, PEX19 N50-His, and StrepII-Hip fusion proteins. The expression plasmids encoding these proteins were transformed into the BL21 *E. coli* strain. LB medium, containing ampicillin, was inoculated with single colonies and incubated overnight at 37°C while shaking. The following day, the cultures were reinoculated at 0.1 OD^600^/mL and further incubated at 37°C with shaking until cell density reached 0.6 OD^600^/mL. Protein expression was induced with 1 mM IPTG for 4 h at 30°C, except for GST-PEX3Δ44, which was induced with 0.4 mM IPTG for 16 h at 16°C. Harvested cell pellets were stored at −20°C before use.

For protein purification, harvested cell pellets were resuspended in PBS with protease inhibitors (5 μg/mL Antipain, 2 μg/mL Aprotinin, 0.35 μg/mL Bestatin, 6 μg/mL Chymostatin, 2.5 μg/mL Leupeptin, 1 μg/mL Pepstatin, 0.1 mM PMSF, 25 μg/mL DNAse, and 1 mM DTT). Cells were disrupted using EmulsiFlex, and unbroken cells were removed by centrifugation at 4,000g (rotor SX4400, Beckman Coulter) for 15 min. The resulting supernatant (SN1) was subjected to high-speed centrifugation at 24,000 g for 1 h (rotor SS-34, Thermo Scientific), yielding supernatant 2 (SN2), a soluble fraction that included overexpressed proteins. Proteins were purified by affinity chromatography using Glutathione Agarose 4B beads for GST-tagged proteins, Protino Ni-NTA Agarose for His-tagged proteins, and Strep-Tactin®XT 4Flow® resin for Strep-tagged proteins. SN2 was incubated with the pre-equilibrated resin for the respective tagged proteins for 2 h in a tube rotator. After collecting the flow-through using a gravity flow column, the protein-bound beads were washed five times with PBS. Proteins were eluted with the appropriate elution buffer: 10 mM reduced glutathione in 50 mM Tris-Cl (pH 8.0) for GST-tagged proteins and 200 mM imidazole in PBS (pH 8.0) for His-tagged proteins. Purification of Strep-tagged proteins was performed according to the manufacturer’s protocol (Iba, cat. no. 2-5010-002). The buffer of the eluted proteins was exchanged to PBS using Amicon centrifugation tubes with a molecular weight cut-off (MWCO) of 10 kDa. The concentration of the proteins was determined by the Bradford method (Thermo, Coomassie Plus assay kit), and protein aliquots were stored at −80°C. All purification steps were performed at 4°C. PBS: 10 mM Na^2^HPO^4^. 2H^2^O, 1.76 mM KH^2^PO^4^ 137 mM NaCl, 2.7 mM KCl, pH 7.4

### *Trypanosoma* cell culture and transfection

In this study, *T. b. brucei* bloodstream form 90-13 (BSF) and procyclic form 29-13 (PCF) cell lines, which co-express T7 RNAP and TetR, were used. PCF cells were grown in SDM-79 medium at 28°C^45^ and the BSF cells were cultured in HMI-11 medium at 37°C with 5% CO_2_^46^. Both cell lines were supplemented with heat inactivated 10% fetal bovine serum. The PCF cultures were maintained at a density of 1 × 10^6^ to 30 × 10^6^ cells/mL, while the BSF cultures were kept in the logarithmic growth phase (cell density below 2×10^6^ cells/mL). Plasmid constructs linearized with the NotI restriction enzyme were transfected into cells as described previously^47^, to integrate stably into the ribosomal RNA locus and the resulting clones were selected using the below described antibiotics. for PCF clones: 10 μg/ml blasticidin for pGN1/pGC1, or 5 μg/ml phleomycin for p2T7-177. For BSF clones: 5 μg/ml blasticidin for pGN1/pGC1, or 2.5 μg/ml phleomycin for p2T7-177.

### Affinity isolation of PEX19 interactome using *in vitro* pulldown

The PCF cells were grown to a density of 20 × 10^6^ cells/mL, harvested (∼500 mL), and snap frozen. Frozen cells were thawed, resuspended in PBS containing 1x protease inhibitor cocktail (PIC) and 1 mM EDTA, and permeabilized with 0.08 mg of digitonin/mg of protein. After a 5- min incubation at room temperature, the cell suspensions were centrifuged at 24,000 g (rotor SS-34, Thermo Scientific) for 30 min to obtain the cytosol-enriched fraction. The sedimented organellar pellet was then treated with a higher digitonin concentration of 3 mg/mg of protein and incubated for 30 min at 4°C on a Mini Rocker. Finally, the samples were centrifuged at 24,000g (rotor SS-34, Thermo Scientific) for 30 min to yield the solubilized organelle-enriched fractions.

Concurrently, 350 µg of recombinant proteins, including GST-PEX19FL, GST-PEX19 ΔΝ30, and GST as a negative control, were allowed to bind to 100 µL of settled glutathione agarose beads by gently rotating the tubes at 4°C for 2 h. After incubation, the beads were washed with PBS to remove unbound proteins. The *Trypanosoma* cytosol or organelle enriched fractions, prepared using the digitonin treatment, were then added to the glutathione agarose beads pre-bound with the recombinant proteins in separate tubes as follows: 1. GST alone, 2. GST + Cytosol fraction, 3. GST + organelle fraction, 4. GST-PEX19^FL^ alone, 5. GST-PEX19^FL^ + Cytosol fraction, 6. GST-PEX19^FL^ + organelle fraction, 7. GST-PEX19^ΔΝ30^ alone, 8. GST- PEX19^ΔΝ30^ + Cytosol fraction, 9. GST-PEX19^ΔΝ30^ + organelle fraction. The tubes were then gently rotated at 4°C for 2 h to facilitate the binding of *Trypanosoma* proteins to GST-PEX19^FL^ and GST-PEX19^ΔN30^. Prior to elution, the beads were washed five times with PBS, with each wash performed by centrifugation to remove unbound proteins. Specifically bound proteins were eluted using 2 units of thrombin in 120 µL PBS, with incubation at 16°C for 16 h to ensure efficient thrombin cleavage. Finally, the eluted samples were analyzed by SDS-PAGE analysis followed by silver staining (described in^48^) and subjected to proteomic analysis to identify the bound proteins.

### Proteolytic in-gel digestion

Thrombin-eluted complexes prepared as described above in triplicates were processed for tryptic in-gel digestion as described previously^49^. Approximately 35 µL of each eluate was separated on 14% Tris-glycine Novex Wedgewell denaturing gels (Invitrogen) and visualized using colloidal Coomassie Brilliant Blue. Each gel lane was cut into 10 slices. Following destaining, gel slices were treated with 5 mM Tris(2-carboxy-ethyl) phosphine prepared in 10 mM ammonium bicarbonate (ABC) to reduce cysteine residues (30 min at 37 °C). Free thiol groups were then alkylated with 50 mM chloroacetamide/10 mM ABC (30 min at room temperature). In-gel digestion was carried out overnight at 37 °C using 0.06 μg of sequencing-grade trypsin (Promega) dissolved in 10 mM ABC per gel slice. Resulting peptides were extracted by two successive rounds of incubation in 0.05% (v/v) trifluoroacetic acid (TFA) and 50% (v/v) acetonitrile (ACN) using an ultrasonic bath (10 min at 4 °C each). Corresponding peptide-containing supernatants were pooled, dried under vacuum, and subsequently desalted with StageTips^48^. To this end, StageTips were conditioned sequentially with methanol, 80% ACN in 0.5% acetic acid (v/v), and 0.5% (v/v) acetic acid. Peptides were loaded onto StageTips, washed twice with 0.5% (v/v) acetic acid, and eluted with 80% ACN/0.5% acetic acid (v/v). Peptides were dried and stored at −80 °C until further analysis.

### Liquid chromatography-mass spectrometry analysis

Dried peptide mixtures were resuspended in 0.1% TFA and analyzed by nano high-performance liquid chromatography-electrospray ionization-tandem mass spectrometry (Nano-HPLC-ESI-MS/MS) using an Orbitrap Elite hybrid mass spectrometer (Thermo Fisher Scientific, Bremen, Germany) coupled to an UltiMate 3000 RSLCnano HPLC system (Thermo Fisher Scientific, Dreieich, Germany). The RSLC system was operated with C18 pre-columns (nanoEase M/Z Symmetry C18; 20 mm length, 0.18 mm inner diameter) and an analytical C18 reversed-phase nano LC column (nanoEase M/Z HSS C18 T3; 250 mm length, 75 mm inner diameter, 1.8 mm particle size, 100 Å packing density). A binary solvent system was employed for peptide separation, composed of 4% (v/v) dimethyl sulfoxide (DMSO)/0.1% (v/v) formic acid (FA) (solvent A) and 30% (v/v) ACN/48% (v/v) methanol/4% (v/v) DMSO/0.1% (v/v) FA (solvent B). Peptides equivalent to 1 µg of protein were loaded, pre-concentrated and washed on the precolumn for 5 min using solvent A and a flow rate of 10 µL/min. Peptides were eluted using the following gradient: 1–7% solvent B in 5 minutes, 7–65% B in 30 min, 65-80% B in 15 min, and 3 min at 80% B at a flow rate of 300 ml/min. Eluted peptides were directed to a fused silica emitter for electrospray ionization using a Nanospray Flex ion source with a DirectJunction adaptor, applying a spray voltage of 1.8 kV and a capillary temperature of 200°C. Mass spectrometric data were acquired in data-dependent mode using the following parameters: MS precursor scans at *m/z* 370–1700 with a resolution of 120,000 (at *m/z* 400); automatic gain control (AGC) of 1 × 10^6^ ions; a maximum injection time (IT) of 200 ms; a TOP20 method for low-energy collision-induced dissociation of multiply charged precursor ions with a normalized collision energy of 35%, an activation q of 0.25, and an activation time of 10 ms; AGC for MS/MS scans of 5 × 10^3^ ions with a maximum IT of 150 ms; and a dynamic exclusion time of 45 sec.

### MS data analysis

For the analysis of mass spectrometric data, the MaxQuant software package (version 1.6.10.43)^50^ and its integrated search engine Andromeda^51^ were employed. Raw data were searched against all protein sequences of *T. brucei* (strain Lister 427, TREU427) obtained from the TriTrypDB (release 48; https://tritrypdb.org/tritrypdb/app) to receive information about proteins present in the sample. ‘Trypsin/P’ was selected as proteolytic enzyme, allowing up to three missed cleavages, and mass tolerances were set to 20 ppm for precursor ions and 0.5 Da for fragment ions. The options ‘match between runs’ and ‘iBAQ’ (i.e., intensity-based absolute quantitation) were activated. Protein identification was based on the detection of at least one unique peptide of seven or more amino acids, applying a false discovery rate of 0.01 to both peptide and protein identifications. For all other parameters, MaxQuant default settings were used, including the fixed modification of carbamidomethylation for cysteine residues and variable modifications set for N-terminal acetylation and methionine oxidation.

For data analysis and visualization, MaxQuant results were processed using the autoprot Python module (v0.2)^52^. To identify proteins specifically associated with PEX19FL or PEX19ΔΝ30, PEX19FL/GST, PEX19ΔΝ30/GST and PEX19ΔΝ30/PEX19FL protein abundance ratios were calculated based on iBAQ MS intensities, for both cytosolic and organellar fractions. Proteins were required to be identified in at least two out of three replicates per experimental condition. First, iBAQ values were log^2^ transformed. Missing values (i.e., in case a protein was only identified in 2/3 replicates) were imputed by drawing random values from a distribution matching the iBAQ value distribution of the existing values shifted downward by 1.3 standard deviations and scaled to a width of 3%. Log^2^ abundance ratios as indicated above were calculated and normalized between equal conditions using cyclic loess normalization^53^. The rank-sum test implemented in autoprot based on the R package RankProd (version 3.11)^54^ as then used to determine proteins specifically enriched with a given bait protein.See **Table S1** for detailed results. A Jupyter notebook providing documentation of the analysis pipeline and statistical tools used is available at https://github.com/ag-warscheid/Tb_PEX38_manuscript.

### Yeast two-hybrid analysis (Y2H)

Yeast two-hybrid assays, including both a colony-lift filter assay (also referred to as a plate-based assay) and a growth-based assay, were performed to investigate protein-protein interactions. As described in^47^, the colony-lift filter assay was conducted using the *S. cerevisiae* wild-type strain PCY2. The growth-based Y2H assay, as described in^55^, was performed using the *S. cerevisiae* strain PJ69-4A. Yeast cells were transformed using the traditional lithium-acetate method, and the interaction analysis was carried out for both assays as described in^47^ and^55^. The interaction between the GAL4 activation domain fused to *Sc*PEX5 and the GAL4 DNA-binding domain fused to *Sc*PCS60 was used as a positive control for the study. Additionally, the GAL4-AD fusion of PEX19, Hip, or *Ld*PEX19, as well as the various GAL4- BD fusions of PEX38 (wild-type or mutants), Hip, or *Ld*PEX38 (wild-type or mutants), were tested for autoactivation, serving as negative controls.

### Peptide array

To identify the binding sites and conduct mutational analysis, *Tb*PEX19 and *Tb*PEX38 peptide arrays were obtained. The immobilized peptides comprised 25 amino acids for PEX19 and 15 amino acids for PEX38, sequentially overlapping by 20 and 13 residues, respectively, to represent the entire sequences of PEX19 and PEX3. These peptides were synthesized on a cellulose membrane, as described previously in^56^. For the proline walk, 15-amino-acid-long peptides, i.e., PEX38 89-103, were used, substituting each residue with a proline. The peptide array was first activated with ethanol for 10 min with gentle shaking, followed by three 10-min washes with TBS. Subsequently, the peptide array was incubated with a blocking buffer (5% fat free milk powder + 0.05% tween 20 + 5% Sucrose in TBS) for 1 h at room temperature. The purified recombinant proteins GST-PEX19, GST-PEX38, or GST alone with a final 1 µM (15 mL) concentration were incubated with the protein arrays for 1 hour at 4°C. Subsequently, the arrays were subjected to three 10-min washes in a TBS buffer at room temperature. This was followed by a ∼16 h incubation with an anti-GST (Sigma, 1:2,000 in blocking buffer) monoclonal antibody at 4°C. After three more TBS washes, a secondary antibody (Horseradish peroxidase-coupled anti-mouse IgGs, 1:5,000 in blocking buffer) was applied, and the arrays were incubated for 1 h at room temperature. Finally, the arrays were scanned using a chemiluminescence substrate (WesternBright Sirius) and the Azure Sapphire biomolecular imager. TBS: 50 mM Tris, 137 mM NaCl, 2.7 mM KCl, pH 8.

### *In vitro* pull-down assay of PEX19 N50-His^6^

For the *in vitro* pull-down assay, 10 µL of settled glutathione agarose beads were separately incubated with 100 µg of recombinant GST and GST-PEX38 proteins. The incubation was carried out for 2 h at 4°C with gentle rotation. After washing the beads three times with TBS to remove unbound proteins, 20 µg of recombinant PEX19 N50-His^6^ containing 5% sucrose was added to the beads. This allowed the PEX19 protein to bind to either the GST-PEX38 or the control GST for an additional 2 h at 4°C with gentle rotation. Following three more TBS washes, the bound proteins were eluted with 40 µL of 10 mM reduced glutathione in 50 mM Tris, pH 8. The eluted samples were analysed using SDS-PAGE followed by Coomassie staining and immunoblotting.

### Cleavage of GST-tag with biotinylated thrombin

To obtain tag-free proteins for the displacement/competition experiments, 2.5 mg of GST- PEX38 (full-length) was incubated with 1.5 units of biotinylated thrombin (69672-50UN, Sigma-Aldrich) in 1 mL of PBS at 4°C for ∼16 h. The biotinylated thrombin was then captured using a 50 µL magnetic slurry of Dynabeads™ MyOne™ Streptavidin T1 (65601, Invitrogen).

After a 60-min incubation at 4°C with gentle rotation, the samples in the microfuge tube were magnetically separated by placing the tube in a magnet rack for 2-3 min. The tube containing a mixture of the cleaved GST tag and the tag-free protein was carefully decanted. To remove the cleaved GST tag from the tag-free proteins of interest, 350 µL of settled glutathione agarose beads were added and incubated at 4°C for 2 h. Following the incubation, the flow-through containing the tag-free protein was collected. The beads were either denatured or eluted with 10 mM reduced glutathione in 50 mM Tris. The samples collected from each step were analyzed using SDS-PAGE and Coomassie staining to assess the cleavage efficiency and protein purity. In addition, the same tag-free PEX38 65-134 that was used for the structural study was also used for the competition assay.

### AlphaScreen-based assay

The AlphaScreen-based assay was employed for the binding study, competition, or displacement assay, as well as to assess the formation of the ternary complex. To investigate the interaction between the C-terminal His6-tagged PEX19 (full-length and N50) and N-terminal GST-tagged PEX38 (full-length and 69-133), the proteins were used at a concentration of 30 nM in the AlphaScreen system. The protein solution was prepared in PBS (pH 7.4) with 0.5% BSA, and the donor and acceptor beads were prepared in the same buffer with 0.05% Tween 80, both at a concentration of 5 μg/mL. The bead information and assay details were as described previously^47^. The binding assay was performed in three biological replicates, with 6 technical replicates each.

The competition assay was designed to demonstrate that untagged PEX38 (full-length or 65-134) can displace GST-tagged PEX3Δ44 from binding to PEX19-His. Here, 30 nM of PEX19-His was added to a 384-well plate, followed by the addition of 30 nM of GST- PEX3Δ44. Serially diluted untagged proteins, either full-length PEX38 or the 65-134 amino acid region of PEX38, were then added. The dilution was a 2-fold series with 12 points, starting from 0 to 3660 nM for full-length PEX38 and 0 to 5000 nM for the 65-134 amino acid region. The bead concentration and buffer were as described above. The assay was performed in three biological replicates, with 3 technical replicates each.

An AlphaScreen-based complex assay was performed to demonstrate the bridging function of PEX38, which results in the formation of a ternary complex involving PEX19, PEX38, and Hip. In this assay, 5 μL of 100 nM PEX19-His was added to a 384-well plate, followed by the addition of 5 μL of serially diluted untagged proteins, either GST or GST- PEX38. The dilution series consisted of 5 points, ranging from 0 to 400 nM for both proteins, with GST serving as a control. Following a 15-min incubation, 5 μL of 100 nM Strep-Hip was added to the pre-incubated mixture and incubated for an additional 30 min. Subsequently, 5 μL of AlphaScreen Nickel-chelate acceptor beads (cat. no. 6760619C, revvity) were added to the mixture, followed by a 15-min incubation at room temperature. Finally, 5 μL of AlphaScreen Strep-Tactin donor beads (cat. no. AS106D, revvity) were added, and the complete 25 μL reaction solutions were incubated for 45 min at room temperature in the dark. The Alpha signals were captured using a Cytation 5 plate reader with a gain value of 180. The binding assay was performed in three biological replicates, with 6 technical replicates each, to ensure the robustness and reliability of the results.

### Protein sample preparation for structural characterization

PEX constructs were transformed into *E. coli* BL21 (DE3) cells and expressed in LB or isotope-enriched M9 minimal medium. Uniformly ^15^N or ^15^N, ^13^C labeled proteins were expressed in H^2^O M9 minimal medium supplemented with 50 µg/mL kanamycin, 1 g/L [U-^15^N] ammonium chloride and 2 g/L hydrated [U-^13^C] glucose as the sole sources of nitrogen and carbon, respectively. After transformation, single colonies were picked randomly and cultured in the medium of choice overnight at 37°C. The next day, cultures were diluted to an optical density of 600 nm (OD^600^) of 0.1 and grown up to a OD^600^ of 0.4-0.6. Protein expression was induced with 0.5 mM IPTG and was carried out for 4 h at 37°C.

The cells were harvested by centrifugation at 6,000 g for 20 min at 4°C. For protein purification the cell pellets were resuspended in lysis buffer (50 mM Tris pH 7.5, 300 mM NaCl, 20 mM imidazole) substituted with lysozyme (from chicken), DNAse and protease inhibitor mix (Serva, Heidelberg, Germany) and lysed by pulsed sonication (10 min, 40% power, large probe, Fisher Scientific model 550) followed by centrifugation at 38,000 g for 45 min. All proteins were purified using gravity flow Ni-NTA (Qiagen, Monheim, Germany) affinity chromatography. The supernatant of the lysate was incubated with Ni-NTA beads (2 mL/1 L culture) for 20 min at 4°C, while rotating. Subsequently, the protein-bound beads were washed with 7 column volumes (CV) high salt buffer (50mM Tris pH 7.5, 750 mM NaCl, 20 mM imidazole) and 10 CV wash buffer (50 mM Tris pH 7.5, 300 mM NaCl, 20 mM imidazole). The elution was performed with 3-5 CV elution buffer (50 mM Tris pH 7.5, 300 mM NaCl, 500 mM imidazole). Dialysis and SUMO cleavage were executed overnight at 4°C in 20 mM Tris pH 7.5, 150 mM NaCl. Further purification was performed with a reverse Ni-NTA column where the flow through containing the cleaved protein of interest was collected and concentrated for size exclusion chromatography using a Superdex S75, 16/600 (Cytiva, Marlborough, US).

### NMR spectroscopy and structure calculation

NMR data were collected using Bruker Avance III or Avance NEO spectrometers operating at 900, 950, or 1200 MHz, equipped with cryogenic probes. All NMR experiments were conducted in NMR buffer (20 mM NaP, pH 6.5, 100 mM NaCl, 1 mM DTT, and 10% D2O) at 298 K in a 5 mm diameter tube. All NMR spectra were processed using Topspin (Bruker Biospin, Rheinstetten, Germany) or NMRPipe and analyzed using CcpNMR Analysis 2.4.2^57^.

The backbone resonances of ^15^N and ^13^C labeled *Tb*PEX38 (65-134) and *Tb*PEX19 (1- 50-SGGY) were assigned based on heteronuclear 2D and 3D experiments, including ^1^H-^15^N- HSQC, HNCA, HN(CO)CA, CBCA(CO)NH, HNCACB, HNCO, HN(CA)CO^58^. ^1^H and ^13^C side chain assignments were obtained using ^1^H-^13^C-HSQC constant time and carbon, as well as proton-evolved HCC(CO)NH^58^ and HCCH-TOCSY (hcchdigp3d, Bruker) experiments. The peptide backbone of free *Tb*PEX38 (65-134) and *Tb*PEX19 (1-50-SGGY) were assigned at concentrations of 360 µM and 380 µM, respectively. Backbone and side chain assignments of double-labeled *Tb*PEX38 (65-134) and *Tb*PEX19 (1-50-SGGY) complexed with non-labeled *Tb*PEX19 (1-50-SGGY) and *Tb*PEX38 (65-134) were obtained at concentrations of 400 µM for the double-labeled and 800-1000 µM for the non-labeled protein.

{^1^H}-^15^N heteronuclear NOE (hetNOE) experiments were performed using the pulse sequence hsqcnoef3gpsi (Bruker, Avance version 12.01.11) with a 4.5 s interscan delay, under the same conditions used for backbone and side chain assignments. NOE values are reported as the ratio of peak heights in experiments with and without proton saturation (hetNOE = I^sat^/I^0^).

Binding studies via NMR titration experiments were conducted at a reference protein concentration of 100 µM. Each titration point was prepared as an individual sample to avoid dilution effects. The protein ligands were added in increasing concentrations up to an 8-fold excess. The chemical shift perturbation (Δδ*^avg^*) was calculated by using formula Δδ*^avg^*=[(Δδ*_H_*)^2^+(Δδ*_N_*\*0.159)^2^]

Inter- and Intradistance restraints were obtained with ^15^N and ^13^C edited 3D NOESY experiments^58^, respectively. Automated peak picking of NOESY spectra was performed with the program Artina^59^ and peak lists were manually cleaned from artefacts using the CcpNMR Analysis 2.4.2 software package^57^. Resonance assignments, TALOS angle restraints^60^ and cleaned NOESY peak lists from all four samples were combined and used as input for a structure calculation with Cyana (version 3.98.15;^61^) leading to an automated NOESY peak list assignment of 90+ %. A total of 2757 unambiguous NOE distance restraints (**for details, see Table S3**) were detected and used to calculate a bundle of 100 conformers, from which the 20 with the lowest Cyana target function were selected for refinement in implicit water in the program amber20^62^ and used to represent the structure ensemble.

### Isothermal Titration Calorimetry (ITC)

Isothermal titration calorimetry (ITC) measurements of 40 µM *Tb*PEX38 with 440µM *Tb*PEX19 were performed as triplicates at 25°C using a MicroCal PEAQ-ITC (Malvern Instruments Ltd. U.K) calorimeter. Buffer conditions were 20 mM Tris pH 7.5, 50 mM NaCl. For all titrations, a titrant dilution control experiment was performed and subtracted before the data were fitted to a one-site binding model using the Malvern Analysis software.

### Subcellular fractionation by density gradient centrifugation

Subcellular fractionation by density gradient centrifugation was employed to analyze the distribution of PEX38 and other organellar markers across the gradient’s fractions in Procyclic *Trypanosoma* PCF cells, as previously described in^33^. The cells were ruptured with silicon carbide in a pre-cooled mortar, followed by differential centrifugation, which yielded a cytosol-enriched fraction and an organelle-enriched pellet. Density gradient centrifugation was then applied to further investigate the distribution of PEX38 within the subcellular compartments.

### Subcellular fractionation using digitonin treatment

Subcellular fractionation was performed on *Trypanosoma* PCF cells expressing GFP-tagged constructs, including GFP, GFP-PTS1, GFP-PEX38, and PEX38-GFP, after induction with tetracycline for approximately ∼24 h. The cells expressing the proteins of interest were treated with digitonin at a final concentration of 0.1 mg/mg of protein for 3-5 min at 37°C, selectively permeabilizing the plasma membrane. This was followed by centrifugation at 20,800 g (Rotor F45-24-11, Eppendorf) for 15 min at 4°C, which separated the samples into a cytosol-enriched supernatant and an organellar pellet. The samples collected from each step were then analyzed using immunoblotting.

### RNA interference and digitonin fractionation

The double-stranded stem-loop PEX38 RNAi construct was cloned into a *Trypanosoma* expression plasmid p2T7-177, as described in the ’**cloning**’ section. This construct was then genomically integrated into the BSF or PCF *Trypanosoma* cells via transfection, as outlined in the **’*Trypanosoma* cell culture and transfection**’ section. Positive clones were subjected to a survival analysis. RNAi was induced by the addition of tetracycline to the transfected *Trypanosoma* cells, which were cultured at a density of 2 × 10^5^ cells/mL for BSF and 1 × 10^6^ cells/mL for PCF. DMSO-treated cells served as negative controls. After 24 h of incubation, the cell counts were recorded, and the cells were diluted back to the initial densities with the addition of tetracycline. Cultures with densities below the initial values were incubated further without dilution or additional tetracycline. The growth of the transfected cell lines treated with DMSO, or tetracycline, was monitored for 4 days in the case of BSF and 9 days for PCF. The experiment was performed in three biological replicates, with two technical replicates for BSF and PCF. The cumulative growth curves were plotted on a logarithmic scale using GraphPad Prism 10.

Biochemical fractionation using digitonin was performed on day 2 RNAi-harvested samples, including both DMSO-treated and RNAi-induced cells, as described previously^33^. The resulting supernatant was analyzed by immunoblotting using various antibodies, including markers such as enolase, GAPDH, phosphofructokinase, glycerol-3-phosphate dehydrogenases (GPD), aldolase, hexokinase, and MtHsp70.

### Complementation assay

Functional complementation analysis was adapted from^20^. For the *in-cellulo* complementation assay, a codon-exchanged, RNAi-resistant PEX38 gene with an N-terminal 2×FLAG tag was custom-synthesized and obtained from GeneCust, France. Using this as a template, mutants were generated with the primers and strategies described in **Tables S2** and **S3**. These tetracycline-inducible, codon-exchanged constructs, comprising wild-type or mutant 2×FLAG- PEX38, were then transfected into the previously generated BSF PEX38 RNAi cells, as described in the ***Trypanosoma* cell culture and transfection** section. In this system, tetracycline induction results in the depletion of endogenous PEX38 and the simultaneous overexpression of the ectopic, codon-exchanged 2×FLAG-PEX38 protein, either wild-type or mutant. Two controls were used to validate the complementation assay: a negative control transfected with an empty vector and a positive control transfected with the wild-type 2×FLAG- PEX38 construct. Upon tetracycline induction, the negative control cannot restore the growth defect caused by PEX38 RNAi, while the positive control should result in near-normal growth, as PEX38 RNAi is functionally complemented by the codon-exchanged, ectopically expressed wild-type PEX38. Growth analysis was performed as described in the **RNA interference and digitonin fractionation** section, with cells transfected with wild-type and mutant constructs monitored for 3 days. Based on the growth behavior, the essentiality of various PEX38 mutations can be assessed.

### Microscopy

*Trypanosoma* cell lines harboring various tetracycline-inducible GFP-tagged constructs were generated, including GFP, GSP-SKL, GFP-PEX38, and PEX38-GFP. These cell lines were either induced with 1 μg/mL tetracycline or treated with DMSO as a negative control. The cells were then sedimented, fixed in 4% paraformaldehyde in PBS at 4°C for 15 min, and processed for imaging as previously described in^33,47^. The fixed *Trypanosoma* cells were stained with an antibody against the glycosomal marker aldolase (1:500), and the stained cells were visualized and imaged using a Carl Zeiss microscope equipped with Zen 3.6 software. The acquired images were processed using deconvolution and merged. Subsequently, the processed images were analyzed with Zeiss Zen 3.2 software (blue edition).

### Synthesis of cDNA

From the BSF day 1 and day 2 RNAi cells, either induced with 1 μg/mL tetracycline or treated with DMSO as a negative control, cells were harvested and RNA isolated using the NucleoSpin RNA kit (cat no. 740955.50, Macherey-Nagel) following the protocol for RNA purification from cultured cells and tissues. cDNA was prepared using the RevertAid First Strand cDNA Synthesis Kit (cat no. K1621, Thermo Scientific) with 400 ng of total RNA. The semi-quantitative analysis of tubulin (RE7000, RE7001) and PEX38 (RE7528, RE7723) mRNA levels was carried out by routine PCR with the prepared cDNA. The primer sequences used for this analysis are described in **Table S6**.

### Bioinformatics analysis

The protein sequences for Human SGTA, SGTB, ECD/Human suppressor of GCR two, and Yeast (*S. cerevisiae*) Sgt2 were obtained from the UniProt database, with the respective IDs O43765, Q96EQ0, O95905, and Q12118. Additionally, the protein sequence for the Human Hsc70-interacting protein (Hip) was also retrieved from the UniProt database with ID P50502. The protein sequences of the identified PEX38 orthologs from the Trypanosomatid parasites *T. brucei*, *T. cruzi*, *L. donovani*, *L. mexicana*, *L. major*, and *L. infantum* were obtained from the TriTrypDB database with accession IDs Tb927.6.4000/ Tb427_060045000, TcCLB.511737.10, and LdBPK_302740.1, LmxM.29.2740, LmjF.30.2740, and LINF_300032600 respectively. For *Diplonema papillatum*, the sequence was retrieved from GenBank using the corresponding accession number KAJ9472742.1, identified through a BLAST search. The protein sequence of the *T. brucei* Hip was also retrieved from the TriTrypDB database with ID Tb427_030056900. The domain architecture and family of the proteins were predicted using the InterPro Scan tool^63^. The multiple sequence alignment was performed with the Clustal Omega tool^64^ and visualised using Jalview software (v 2.11.0) with a percentage identity colour scheme and a conservation threshold of 30%. A percentage identity and similarity matrix of the proteins was also calculated using the SIAS tool with the BLOSUM62 matrix (http://imed.med.ucm.es/Tools/sias.html).

GET/TRC pathway orthologs were identified using BLAST, InterPro, PantherDB domain searches, and OrthoCML, while structural similarity searches were performed with Foldseek. The protein sequences for Get1 and Get3 were obtained from UniProt or TriTrypDB. Get1 orthologs include *Hs*Get1 (O00258), *Sc*Get1 (P53192), and *Bodo saltans* (BSAL_19620). Get3 orthologs include *Hs*Get3 (O43681), *Sc*Get3 (Q12154), *T.cruzi* (*Tc*Get3, TcCLB.510101.490), *L. donovani* (*Ld*Get3, LdBPK_110710.1.1) and *B. saltans* (*Bs*Get3, BSAL_77145).

### Immunoblotting

The samples or cell lysate to be analyzed were denatured in Laemmli buffer for 5 min at 95 °C and separated by SDS-PAGE. Following gel electrophoresis, immunoblotting was performed as described in^33^ . The primary antibodies used in this study were mouse anti-GFP, FLAG, MtHSP70 and human alpha-tubulin (which shows cross-reactivity with *Trypanosoma* tubulin) and rabbit anti-*Trypanosoma* Aldolase, Enolase, Hexokinase, GAPDH, GIM5, PEX11, PFK, G3PDH/GPD, VDAC and PEX38. The dilutions, buffers, and secondary antibodies are described in^33^, except for the FLAG and PEX38 antibodies. The PEX38 antibody was generated in this study using a label-free PEX38 1-134 antigen and was subsequently affinity-purified by Eurogentec. The FLAG (1:2,000, Sigma-Aldrich) and PEX38 antibodies (1 μg/mL) were prepared in PBS-T (Tween-20 0.05%) buffer with 3% BSA. The immunoblots were scanned using the LI-COR Odyssey Infrared Imaging System and analyzed with Image Studio version 5.2.

### Statistical analysis

Statistical analysis for AlphaScreen results was performed using 2-way ANOVA with Šídák’s multiple comparisons test within each row, comparing columns with respective controls. The analysis was based on the values obtained from three independent biological replicates, each with six technical replicates. For the competition assay, the values were obtained from three independent biological replicates, each with three technical replicates. These values were initially transformed to log values using the function X=Log (X), which were further analyzed by non-linear regression, i.e., curve fitting with the equation for log(inhibitor) vs. response -- Variable slope (four parameters) to determine the IC 50. All the analysis was performed using GraphPad Prism (Version 10.3.0).

## Supplementary Figures

**Figure S1:**
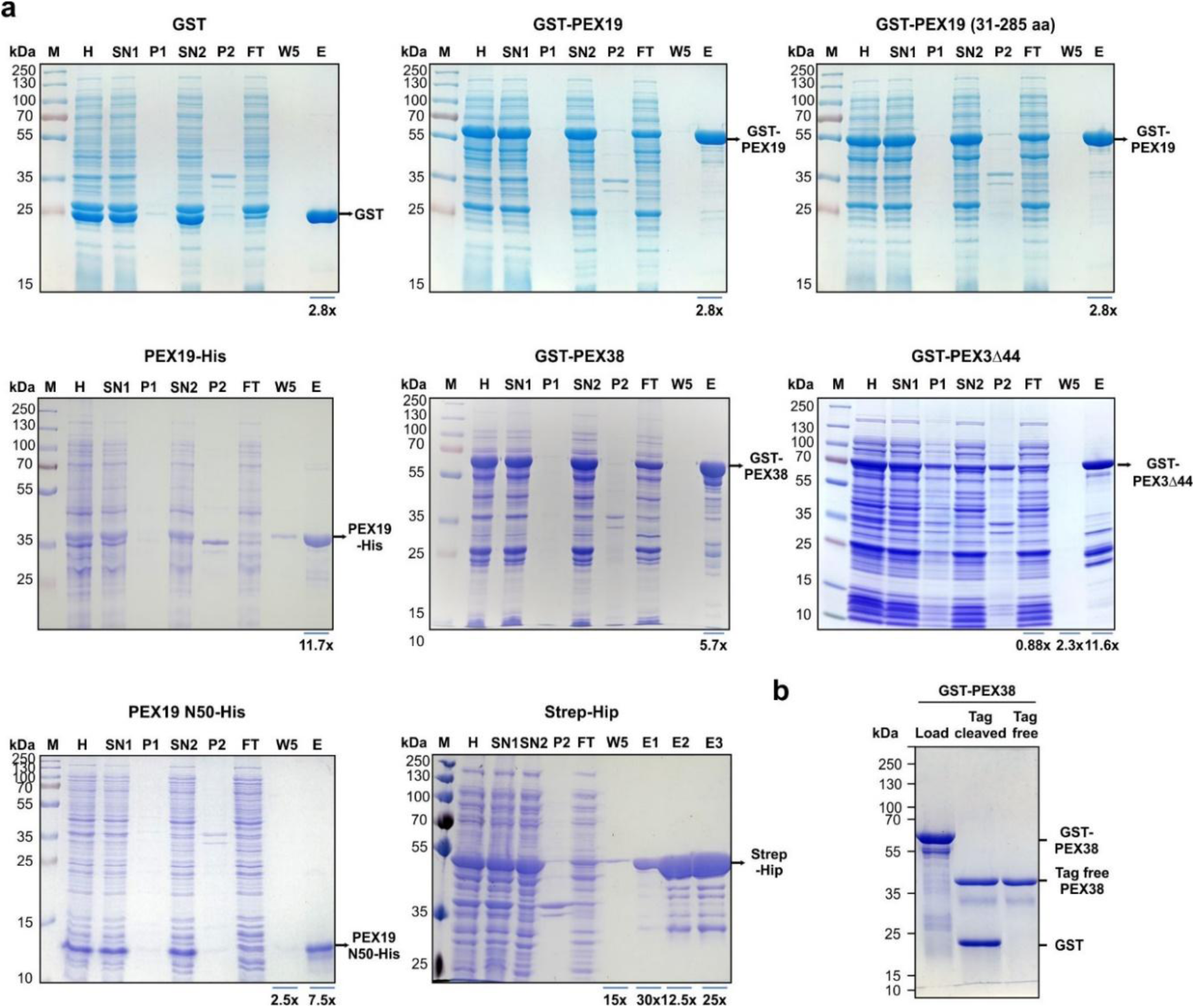
Affinity purification profiles of the recombinant proteins. **a)** Various bacterially expressed proteins were affinity-purified using Glutathione-, Ni-NTA- or Strep-Tactin-XT- Agarose Beads according to the tag as indicated. Samples were collected at each step and analysed by SDS-PAGE and Coomassie staining. The samples include H, Homogenate; SN1, supernatant 1; P1, Pellet 1; SN2, supernatant 2; P1, Pellet 2; FT, flow-through; W5, Wash 5; E, Elution. **b)** Preparation of tag-free PEX38. The GST-tag was cleaved from GST-PEX38 using biotinylated thrombin, which was then captured using streptavidin beads, while the cleaved GST-tag was subsequently removed using glutathione agarose beads.

**Figure S2:**
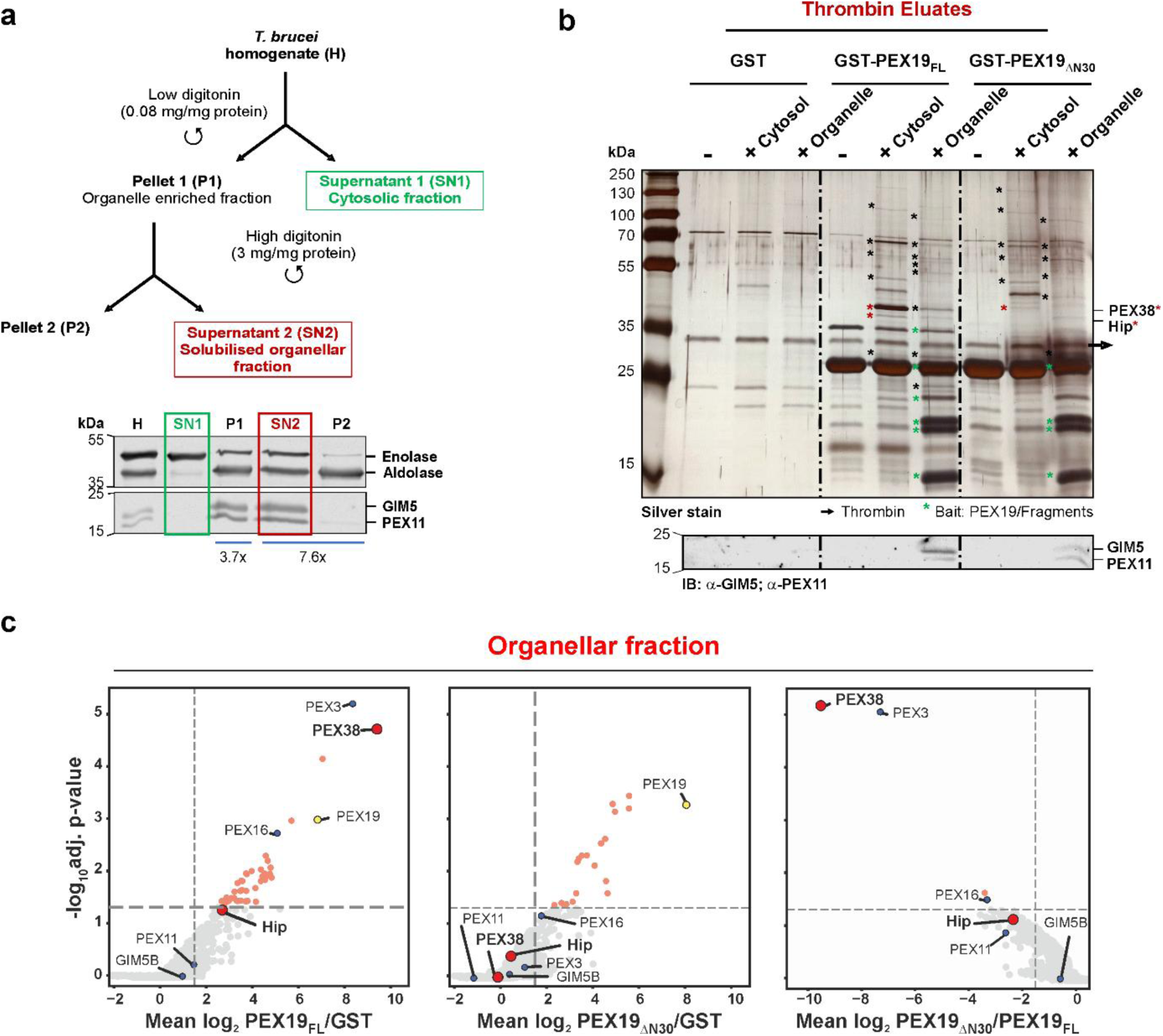
Identification of PEX19-interacting proteins from cytosolic and organellar fractions of trypanosomes: **a)** Scheme for the preparation of cytosol and organelle-rich fractions from procyclic form *Trypanosoma brucei* cells (upper panel). Samples collected at each step of the preparation were analysed by SDS-PAGE followed by immunoblotting with polyclonal antibodies against *T. brucei* enolase, aldolase, GIM5, and PEX11. **b)** Cytosolic and organellar binding partners of *T. brucei* PEX19 were isolated using affinity pull-down using GST alone as control or GST-tagged PEX19 full-length (PEX19FL) or a PEX19 variant lacking the N-terminal 30 amino acids (PEX19ΔN30). The thrombin-eluted fractions were analysed by SDS-PAGE and silver staining. Putative binding partners of PEX19 that are visible as additional bands in both cytosolic and organelle-enriched fractions but not in the control GST are marked with black asterisks. Lower panel: Immunoblot analysis using polyclonal antibodies against GIM5 and PEX11. **c)** Affinity Purification Mass Spectrometry (AP-MS) analysis of the PEX19 (FL and ΔN30) interactomes from trypanosomal organellar fractions. Ratios (left to right) of PEX19FL/GST, PEX19ΔN30/GST , and PEX19ΔN30/ PEX19FL were calculated from quantified iBAQ MS intensities, and a rank-product test was performed for the identification of specifically bound proteins and determination of adjusted p-values ^1^. Highlighted proteins are bait PEX19 (yellow), known glycosomal membrane proteins (blue), PEX38, and HiP (red) and proteins with significant fold change (light red). Vertical dashed lines indicate a log2 ratio of greater than or less than 1.5 (left, middle) or less than 1.5 (right), respectively (n ≥ 2 replicates), and the horizontal line indicates a false discovery rate of 5% (n = 3 biological replicates).

**Figure S3:**
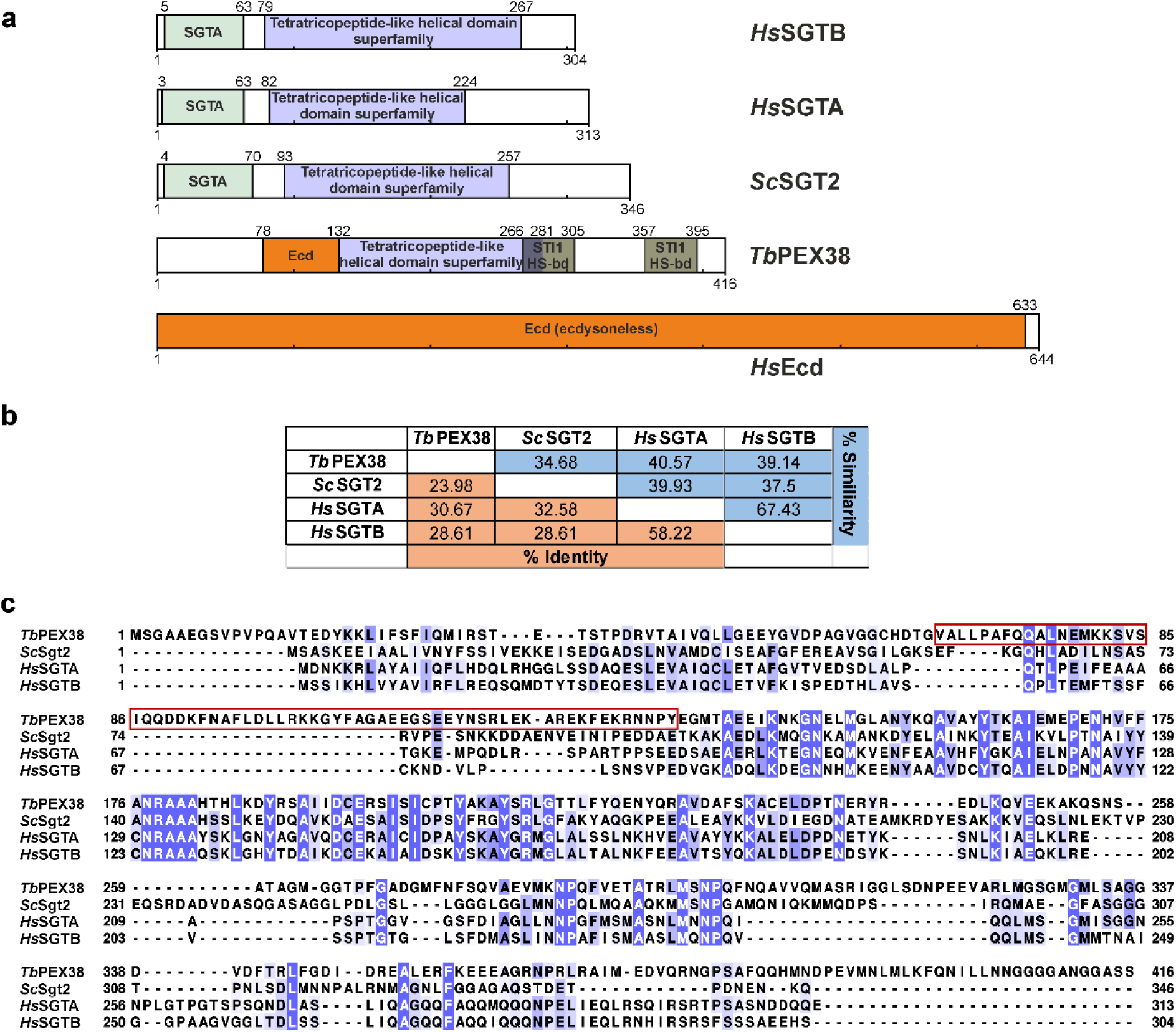
Bioinformatic analysis of PEX38. **a)** Domain architecture of PEX38, human and yeast SGT family proteins and the human Ecd protein. SGTA, SGTA homodimerisation domain; STI1 HS-bd, STress Inducible 1 Heat Shock chaperonin binding domain; Ecd, ecdysoneless, identified using the InterPro domain database. **b)** The percentage identity and similarity matrix of PEX38 with the SGTA or SGTB protein of human and yeast counterparts, generated using SIAS homology modelling. **c)** Multiple sequence alignment of *Sc*Sgt2, *Hs*SGTA/B, and *Tb*PEX38 protein sequences. The sequence conservation is coloured according to the percentage identity with a conservation threshold of 30%. The identified PEX19 binding region within the PEX38 of *T. brucei* is indicated by the red box.

**Figure S4:**
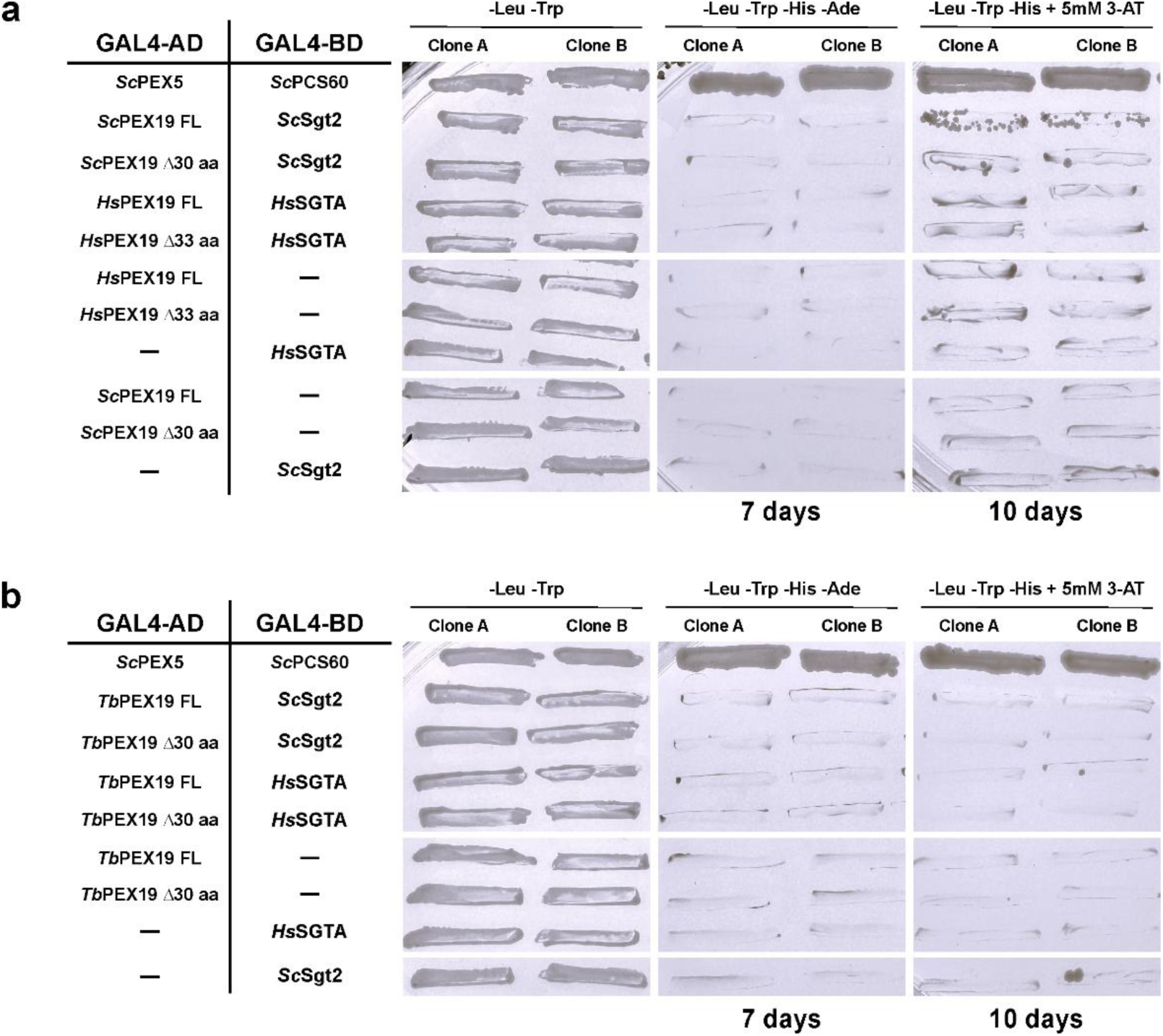
Y2H analysis of Human *(Hs), S. cerevisiae (Sc),* and *T. brucei (Tb)* PEX19 interactions with SGTA and Sgt2. Y2H assays were performed in the *S. cerevisiae* PJ69-4A strain using constructs fused to either the GAL4 activation domain (AD) or DNA-binding domain (BD). **a)** Interactions were tested between full-length (FL) PEX19 and a PEX19 variant lacking the PEX3-binding motif (Δ30aa) with SGTA and Sgt2 from *Hs* and *Sc*, respectively. No interaction was detected between *Hs*PEX19 (FL and Δ30aa) and *Hs*SGTA, except for a very weak interaction observed between *Sc*PEX19 FL and *Sc*Sgt2. **b)** Interactions were also tested between *T. brucei* PEX19 (*Tb*PEX19 FL and Δ30aa) and *Hs*SGTA or *Sc*Sgt2. No interactions were observed for *Tb*PEX19 with SGTA or Sgt2. *Sc*PEX5 and *Sc*PCS60 served as a positive control, while the negative control showed no autoactivation.

**Figure S5:**
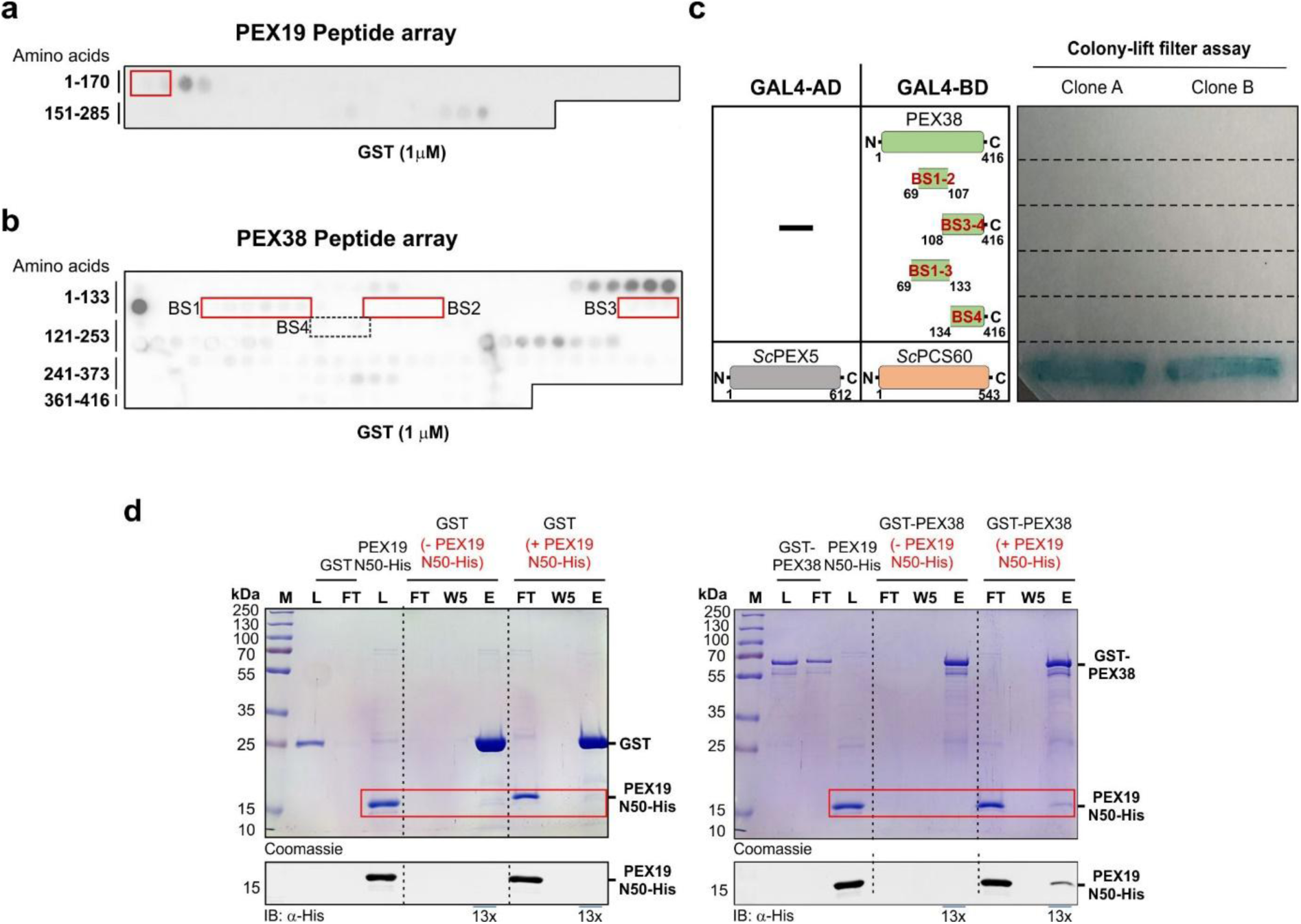
**a, b)** The PEX19 and PEX38 peptide arrays were probed with GST-PEX38 and GST-PEX19 (as shown in Fig. 2a and 2b) as well as with the control protein GST (as shown here), which served as a negative control. Red and black boxes indicate regions with no background signal, while binding was observed with the test proteins. **c)** The GAL4-AD fusion of PEX19 and the various GAL4-BD fusions of PEX38 were tested for autoactivation in Y2H assay. No colour development was observed in the assay, indicating that the constructs do not exhibit autoactivation. *Sc*PEX5-PCS60 served as a positive control for the study (as shown in Fig. 2c). A pull-down assay was performed *in vitro* using recombinant GST-PEX38 (right panel) or GST (left panel, negative control), which were pre-incubated with glutathione agarose beads. This was followed by incubation with the PEX19 N50-His protein. The bound proteins were then eluted using reduced glutathione and analyzed by SDS-PAGE and Coomassie Blue staining. The PEX19 N50-His protein was found to be pulled down with GST-PEX38, which was further confirmed by immunoblotting using an anti-His monoclonal antibody (lower panel).

**Figure S6:**
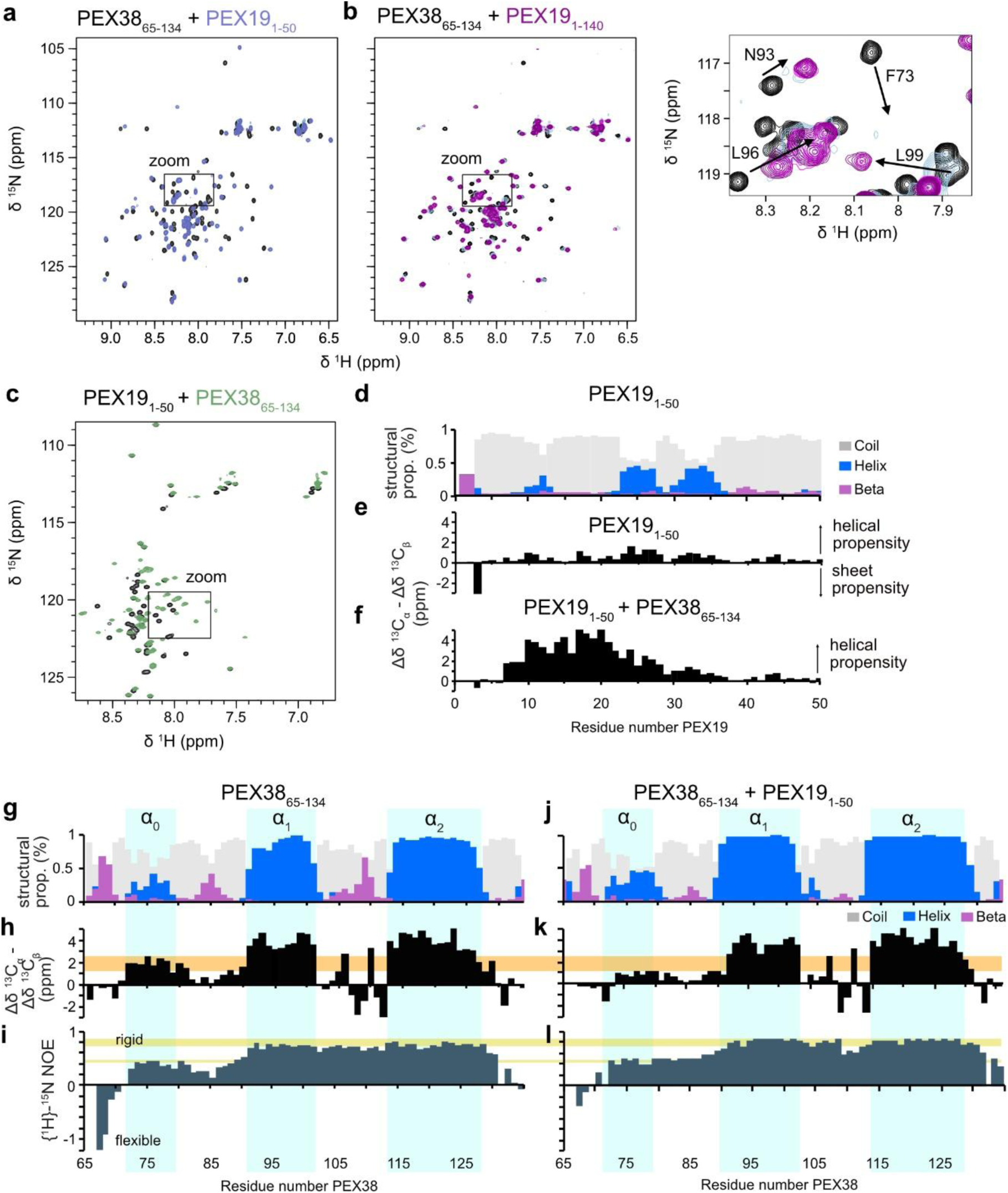
Binding of PEX19 to PEX 38 induces a helix in PEX19 and stabilizes the helical bundle in PEX38. Overlaid ^1^H-^15^N spectra of ^15^N labeled PEX3865-134 (black) titrated with increasing concentrations of **(a)** PEX191- 50 (purple scale; zoom is shown in Fig. 3b) and **(b)** PEX191-140 (magenta scale) with zoom shown on the right. **c)** Overlay of ^1^H-^15^N spectra of PEX191-50 (black) and PEX191-50 saturated with PEX3865-134 (green) (zoomed region is shown in Fig. 3a). **d)** TALOS-N, Secondary structure propensity of free PEX191-50 based on experimentally obtained secondary chemical shifts (Δδ^13^Ca-Δδ^13^Cb) shown in **(e)**. **f**) Experimentally obtained secondary chemical shifts of PEX191-50 bound to PEX3865-134. **g)** TALOS-N Secondary structure propensity, **h)** experimentally obtained secondary chemical shifts and **i)** heteronuclear NOE experiments of free PEX3865-134. **j)** TALOS-N. Secondary structure propensity, **k)** experimentally obtained secondary chemical shifts and **l)** heteronuclear NOE experiments of PEX3865-134 bound to PEX191-50. In **g**) and **j**) propensities for random coil, helix or beta strand secondary structure is shown in gray, blue and purple, respectively. Orange and yellow lines in h) – l) indicate differences in secondary structure and flexibility of free PEX38 (h, j) and PEX38 bound by PEX19.

**Figure S7:**
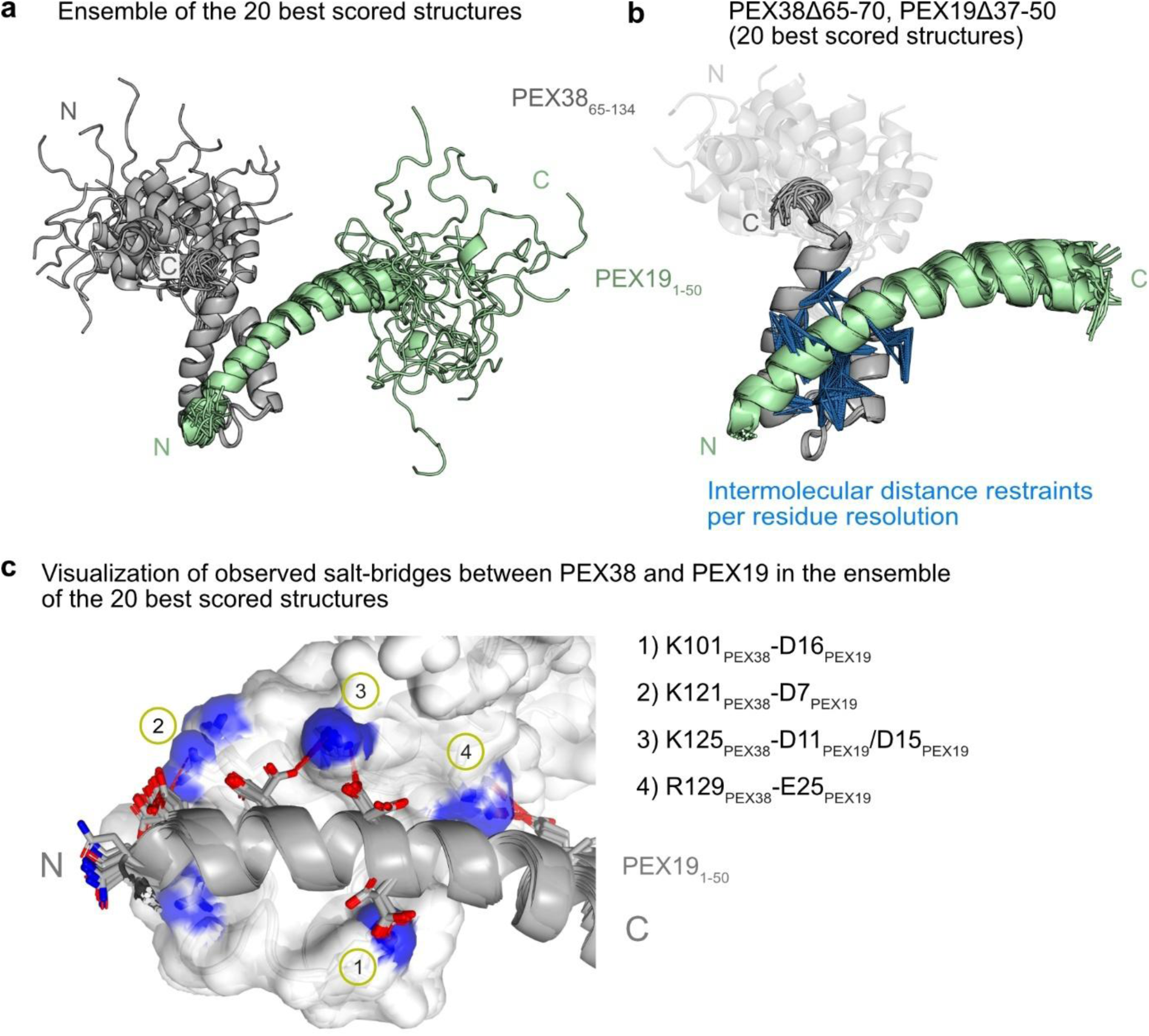
Structure calculation of PEX3865-134 in complex with PEX191-50. **a)** The 20 best scored structures of PEX38 (grey) and PEX19 (green) calculated with CYANA**. b)** Trimmed representation showing the only rigid structures (PEX38Δ65-70 and PEX19Δ37-50) with intermolecular with per residue resolution (not per atom; only one NOE per residue) visualized as blue dashed lines. **c**) Surface representation of PEX39 with cartoon representation of PEX19. Nitrogen atoms of Lys and Arg residues located in the PEX38 binding interface are colored in blue, while oxygen atoms of Asp and Glue residues (shown as sticks) of the PEX19 amphipathic helix are colored in red. Salt-bridges between PEX38 and PEX19 observed in the ensemble were visualized using PyMol 3.1.6.1 (Schrodinger) by detecting distances up to 4 Å between PEX38 Lys NZ or Arg NE and PEX19 Asp OD and Glu OE atoms and are indicated by red dashed lines. A hydrogen bond between PEX38 Arg118 and PEX19 Asn6 was not observed but would be possible as transient contact.

**Figure S8:**
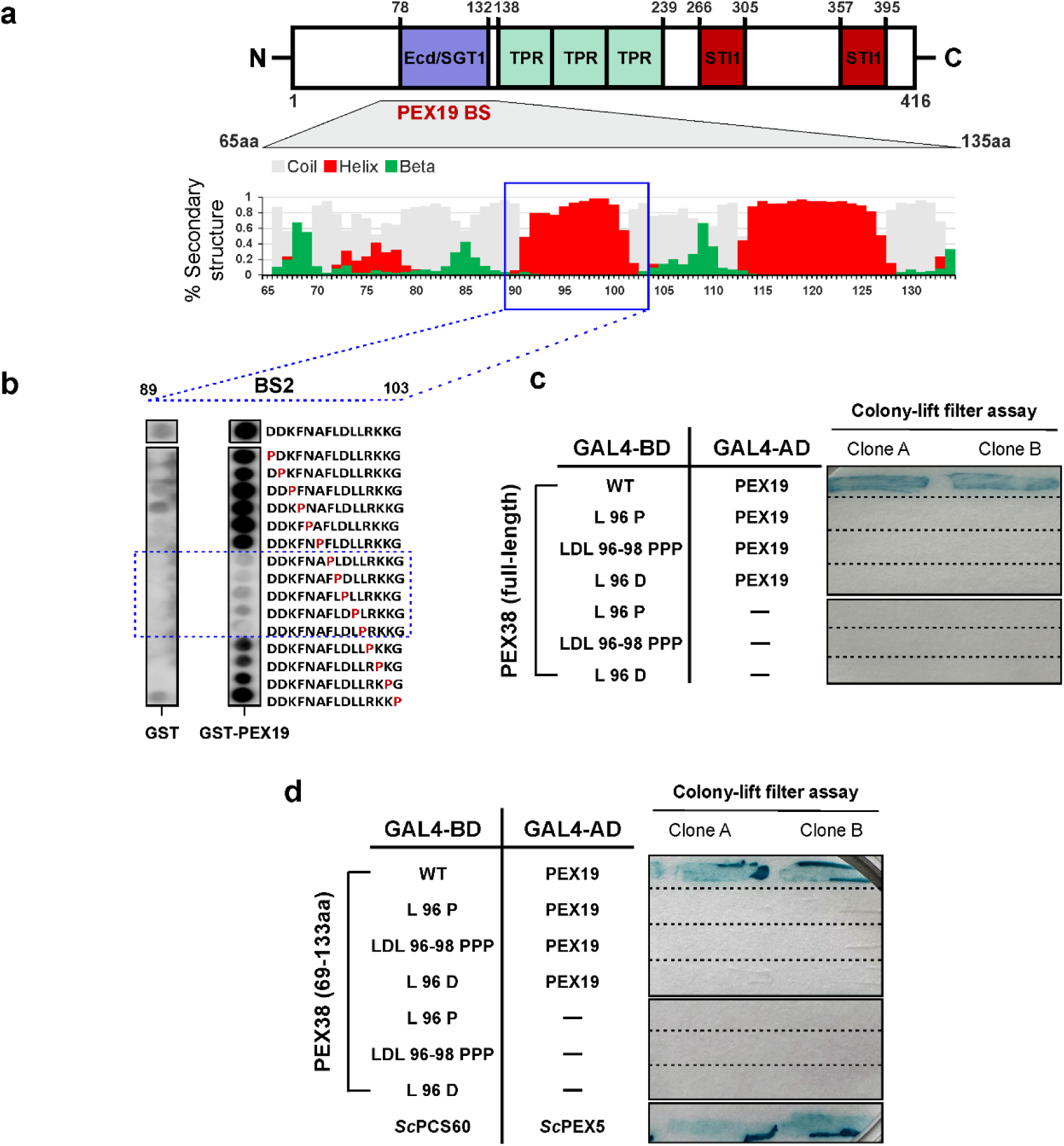
Molecular characterization of the PEX19-PEX38 interaction. **a)** The schematic representation of PEX38 illustrates the domains predicted by the InterPro scan, with the PEX19 binding region highlighted in red. The experimentally determined secondary structure of the PEX19 binding domain (ECD/SGT1) of PEX38 is depicted as structural propensities obtained from secondary chemical shifts (Δδ^13^Ca-Δδ^13^Cb) and TALOS N (**Fig. S6g**). **b)** The identified PEX19 binding region in PEX38 predominantly comprises alpha-helical structures. From this region, the central PEX19 binding site, BS2, which consists of a 15-amino acid peptide, was selected for mutational analysis using a proline walk, as indicated by the blue dotted boxes. Synthetic 15-mer peptides with single proline substitutions at each position within the above-chosen region were tested for interaction with GST as a control or GST-PEX19. The wildtype peptide represents the unaltered sequence, while the proline walk indicates the proline substitution, shown in red font. Within the region spanning amino acids 89-103, replacement of the central residues FLDLL with proline, abolishes their interaction with PEX19, as denoted by the blue dotted box. **c, d)** Yeast two-hybrid (Y2H) assays were performed to investigate the effect of indicated mutations on the interaction between PEX38 and PEX19. Mutating the amino acid residues at positions L96 P/D and LDL 96-98 PPP abolished the interaction of both full-length PEX38 and its 69-133 fragment with PEX19. The assay was performed in triplicate using three different clones.

**Figure S9:**
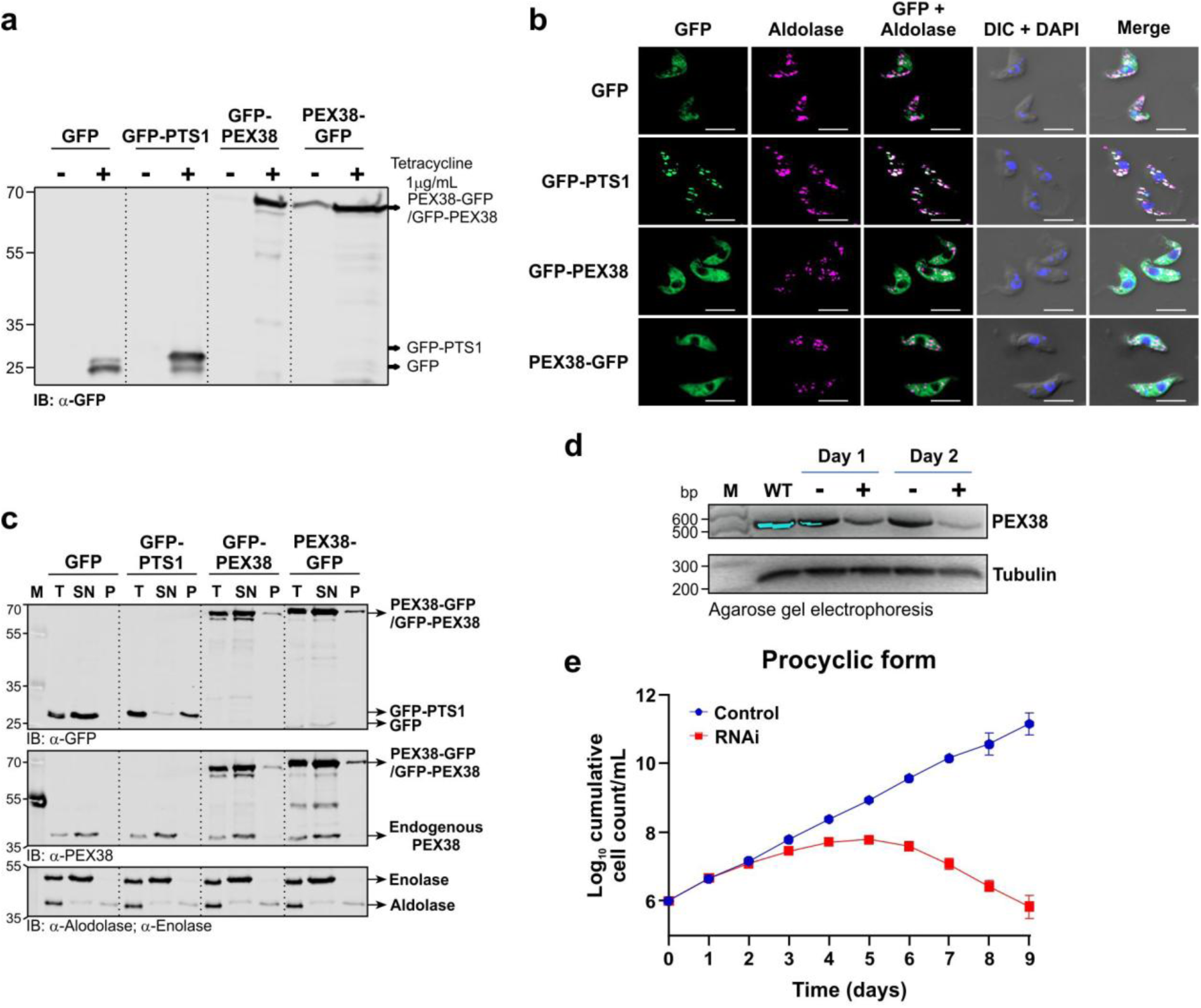
Subcellular localization of GFP-tagged PEX38 proteins in *T. brucei* by fluorescence microscopy and cellular fractionation. **a)** Expression levels with and without tetracycline induction of GFP-fused constructs, including GFP, GFP-PTS1, and PEX38 tagged with GFP at the N- or C-terminus were analyzed by immunoblotting. **b)** Immunofluorescence microscopy of the localization of GFP fusion proteins as well as the glycosomal marker aldolase, and the DAPI-stained nucleus and kinetoplast in the same cell lines. The GFP-PTS1 constructs co-localized with the glycosomal marker aldolase (pseudocolored in magenta). In contrast, the positive control GFP exhibited a cytosolic pattern, as evident from the overall diffuse cell labelling. Similarly, the PEX38 constructs tagged with GFP at the N- or C-terminus also localized to the cytosol. Scale bar – 5 μm. **c)** Cellular fractionation was performed using the same cell lines mentioned above. Immunoblot analysis of fractions was performed using an anti-GFP antibody to detect indicated fusion proteins; enolase and GFP were monitored as cytosolic markers. The lanes represent the following samples: T, total lysate; SN, digitonin supernatant that contains cytosolic proteins; and P, digitonin pellet that contains organellar proteins. **d)** A semi-quantitative analysis of tubulin and PEX38 mRNA levels was performed using routine PCR with cDNA isolated from wild-type and RNAi bloodstream form cells, including uninduced samples and those with RNAi induction on days 1 and 2 (Fig. 4b). **e).** Growth curve demonstrating that PEX38 is an essential protein for the survival of *T. brucei* in the procyclic form.

**Figure S10:**
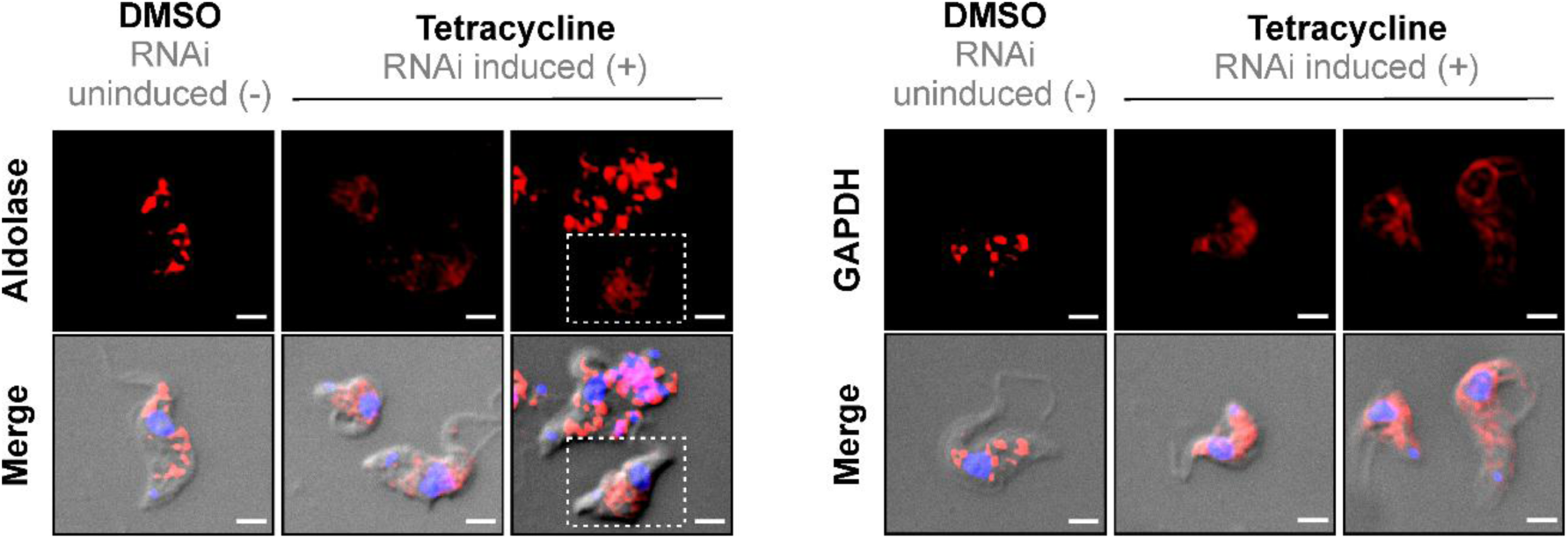
Immunofluorescence microscopy of glycosomes upon PEX38 RNAi. On day 2 of PEX38 RNAi, both DMSO and tetracycline treated cells were analyzed for aldolase and GAPDH by immunofluorescence microscopy. In DMSO-treated cells, aldolase and GAPDH display a punctate glycosomal pattern. Upon PEX38 RNAi induction, glycosomal markers labelling was similar but puncta appeared less bright and more diffuse. Merge channel shows brightfield images with DNA stained by DAPI. The scale bar represents 2 μm.

**Figure S11:**
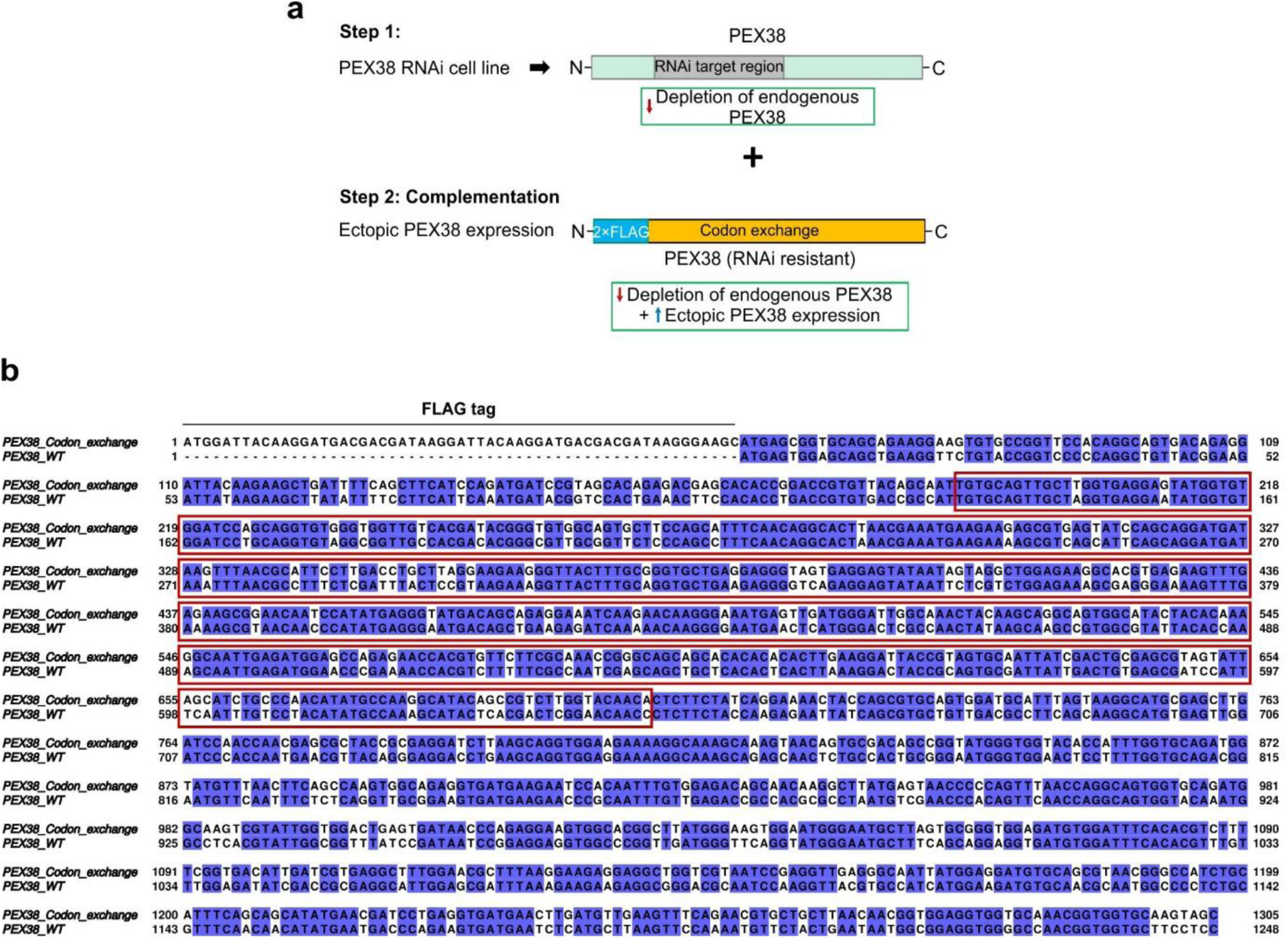
**a)** Steps involved in generating an RNAi-resistant cell line for complementation assay. Step 1: Generation of a PEX38 RNAi cell line by genomic integration of a stem-loop RNAi construct containing the selection marker phleomycin through homologous recombination. This targets the coding region of PEX38 (marked in a grey box) resulting in the depletion of endogenous PEX38 (upper panel). Positive clones, verified by qRT-PCR for the efficient RNAi of PEX38, are further used in step 2 (lower panel). Step 2: Ectopic integration and expression of codon exchanged RNAi resistant 2×FLAG-tagged PEX38 for functional complementation (selection marker blasticidin). Both constructs are integrated into a tetracycline-inducible *Trypanosoma* cell line. Upon tetracycline induction, positive clones exhibit depletion of endogenous PEX38 and overexpression of 2×FLAG-PEX38 protein. **b)** Sequence alignment of wildtype PEX38 gene with a codon-exchanged synthetic gene encoding PEX38. The RNAi target region is marked by a red box. Multiple sequence alignment was performed using the Clustal Omega tool, and the aligned sequences were visualized using Jalview software with the percentage identity colour scheme.

**Figure S12:**
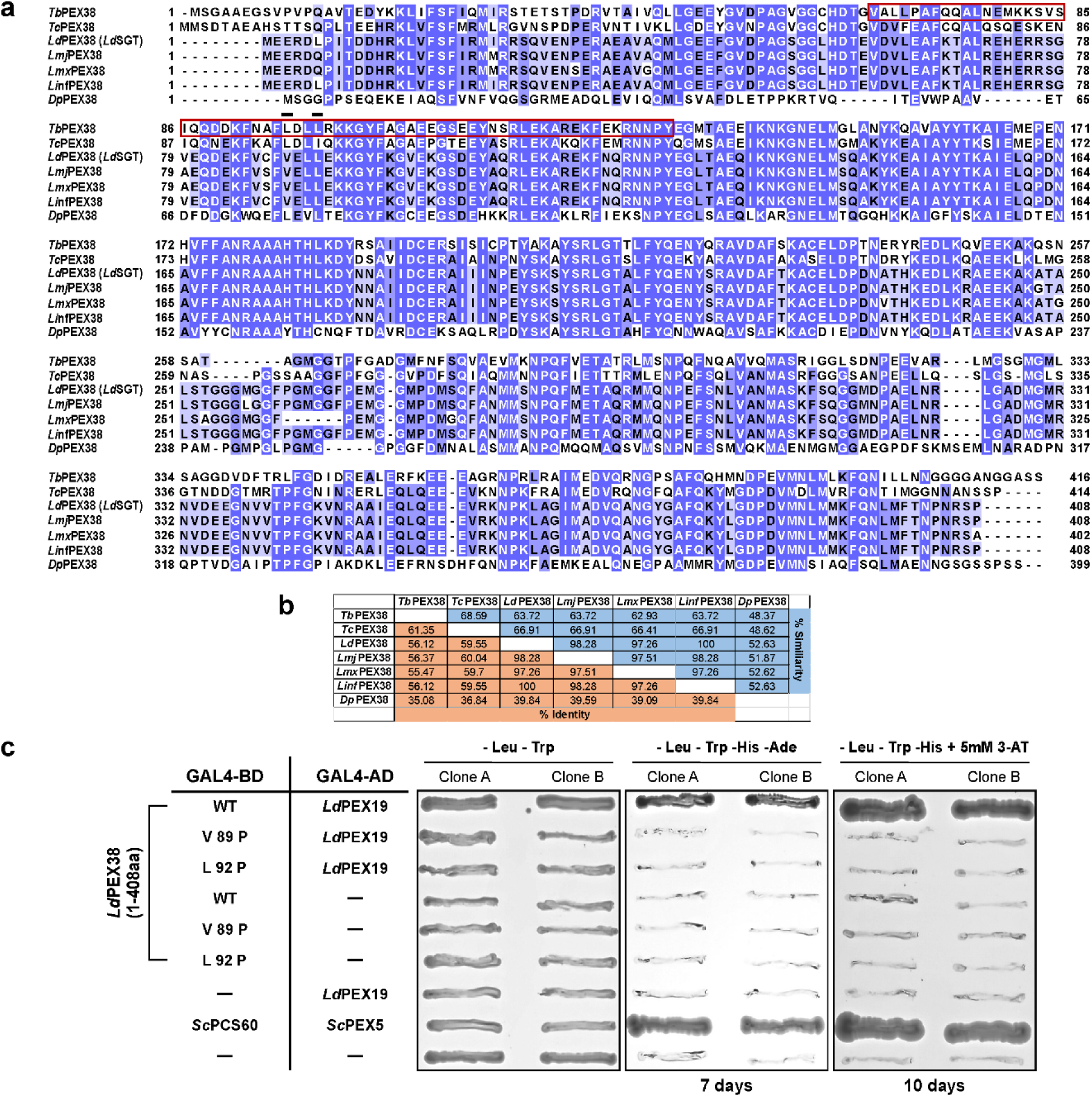
Characterization of the PEX19-PEX38 interaction in clinically relevant *Leishmania parasites.* **a)** Multiple sequence alignment of the PEX38 protein sequences from the trypanosomatid parasites, including *Trypanosoma brucei*, *T. cruzi*, *Leishmania donovani*, *L. infantum*, *L. major*, *L. Mexicana* and *Diplonema papillatum*. Sequence conservation is color-coded based on percentage identity, with a conservation threshold of 30%. The red box highlights the identified PEX19 binding region within *T. brucei* PEX38, while the black lines indicate two conserved residues that are essential for interaction with PEX19. **b)** Analysis of percentage identity and similarity among PEX38 homologs across parasitic organisms using SIAS Homology Modelling. **c)** Yeast two-hybrid (Y2H) interaction analysis of *Ld*PEX38 (wild type or mutant; fused to GAL4 binding domain) and *Ld*PEX19 (fused to GAL4 activation domain) using colony-lift filter assays. The interaction between *Sc*PEX5 and *Sc*PCS60 served as a positive control. The assay was performed in triplicate using three different clones.

**Figure S13:**
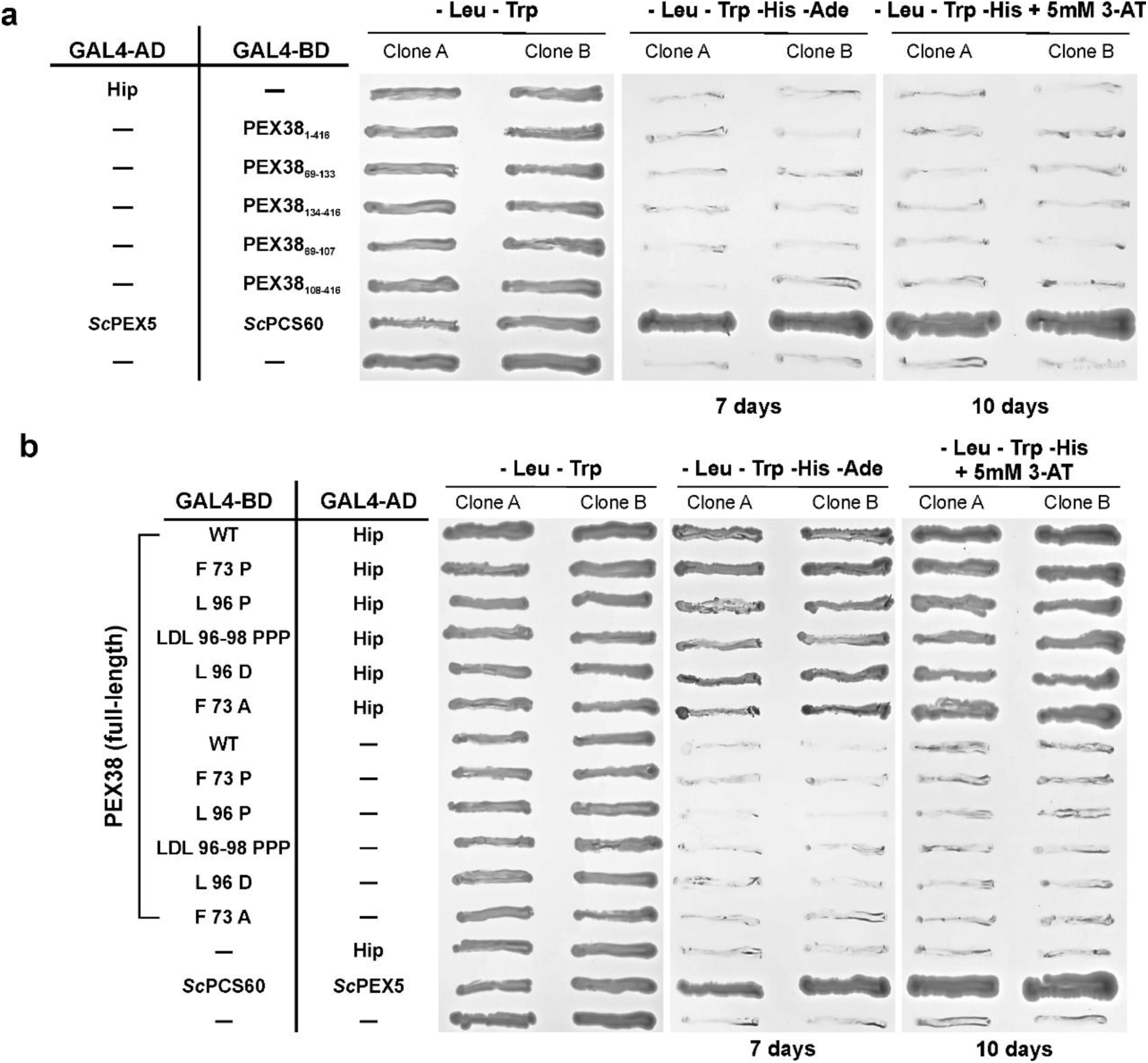
**a)** A Y2H interaction assay was performed between full-length Hip and various PEX38 constructs fused to the GAL4-activation or -binding domain. The constructs were co-transformed into the PJ69-4A yeast strain and analyzed using a growth-based study. These results are negative controls for the study, as shown in Figure 5c. **b)** The effect of PEX38 mutations that disrupt the interaction with PEX19 on the interaction with Hip protein was investigated using Y2H assay. Hip and PEX38 (wild type or mutants) constructs were fused to the GAL4-activation or -binding domain and co-transformed into the PJ69-4A yeast strain. The interactions were then analyzed using a growth-based assay.

**Figure S14:**
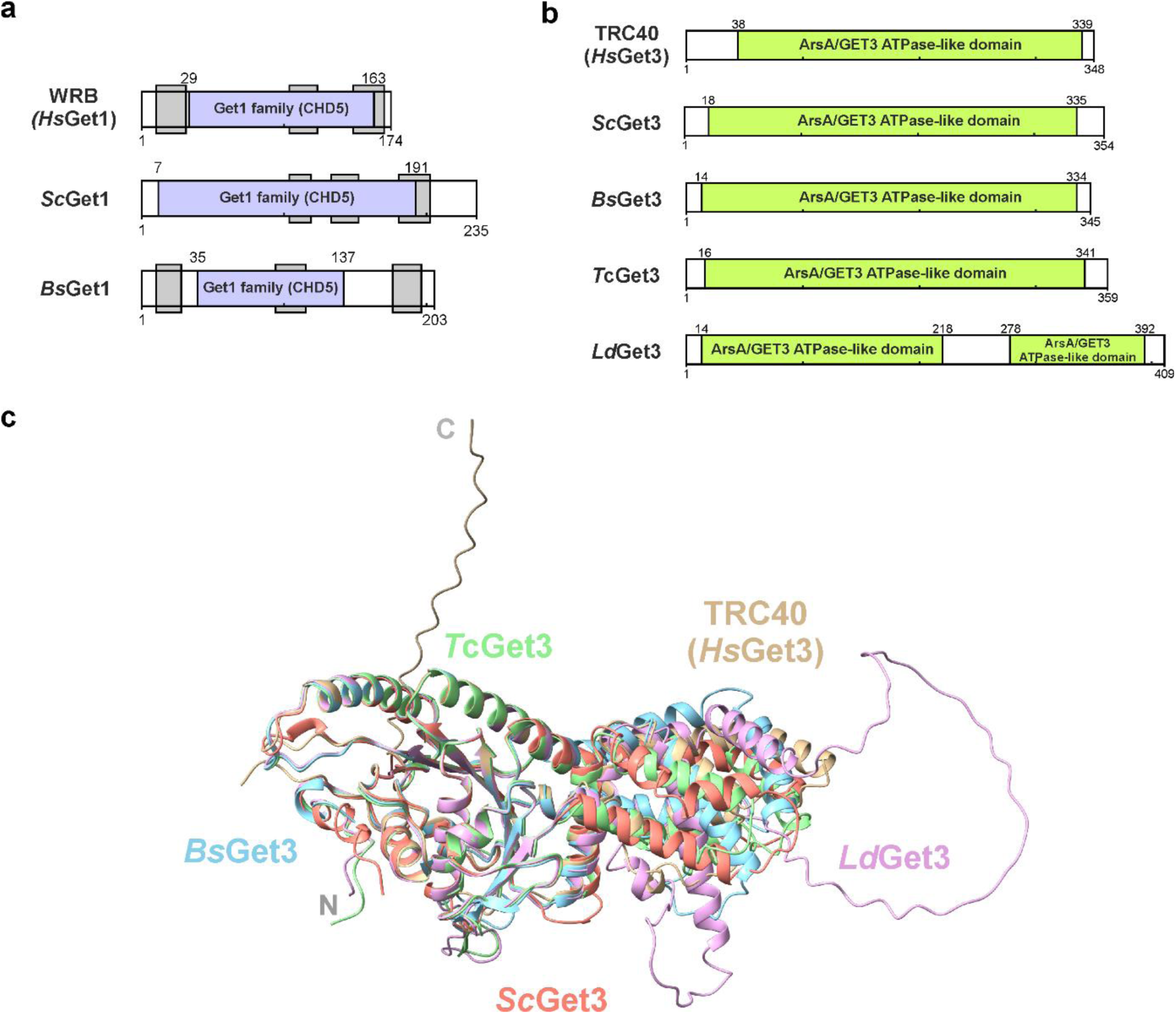
Bioinformatic and structural analysis of Get1 and Get3 orthologs in kinetoplastids. **a)** Domain architecture of Get1 proteins from human *(Hs), S. cerevisiae (Sc),* and *Bodo saltans (Bs)*, revealed by InterPro domain searches. All proteins contain the conserved Get1 family domain (IPR028945), with grey boxes indicating predicted transmembrane segments. b) Domain analysis of Get3 homologs in kinetoplastids (*T. cruzi: Tc*, *L. donovani: Ld*, and *B. saltans: Bs*) alongside *Hs* and *Sc* Get3. All homologs contain the conserved ArsA/GET3 family domain (IPR025723) identified using the InterPro database. c) AlphaFold-predicted structures of kinetoplastid Get3 homologs (*Bs, Tc, Ld*) show high structural similarity to *Hs* and *Sc* Get3. Structural superpositions were performed using *Hs*Get3 as the reference. Colors indicate species: *Hs*Get3 (tan), *Sc*Get3 (salmon), *Bs*Get3 (blue), *Tc*Get3 (pale green), and *Ld*Get3 (plum).

## Supplementary Tables

**Table S2.**
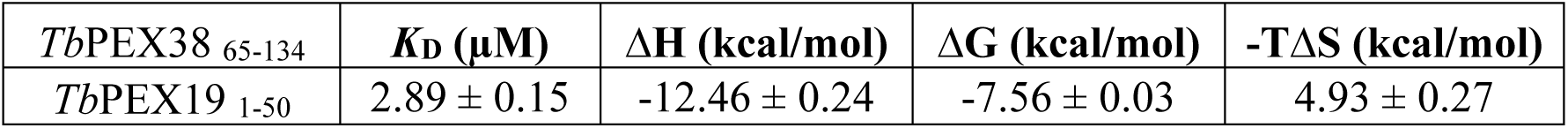
Isothermal titration calorimetry of PEX38^65-134^ titrated with PEX19^1-50^ with n=3.

**Table S3:**
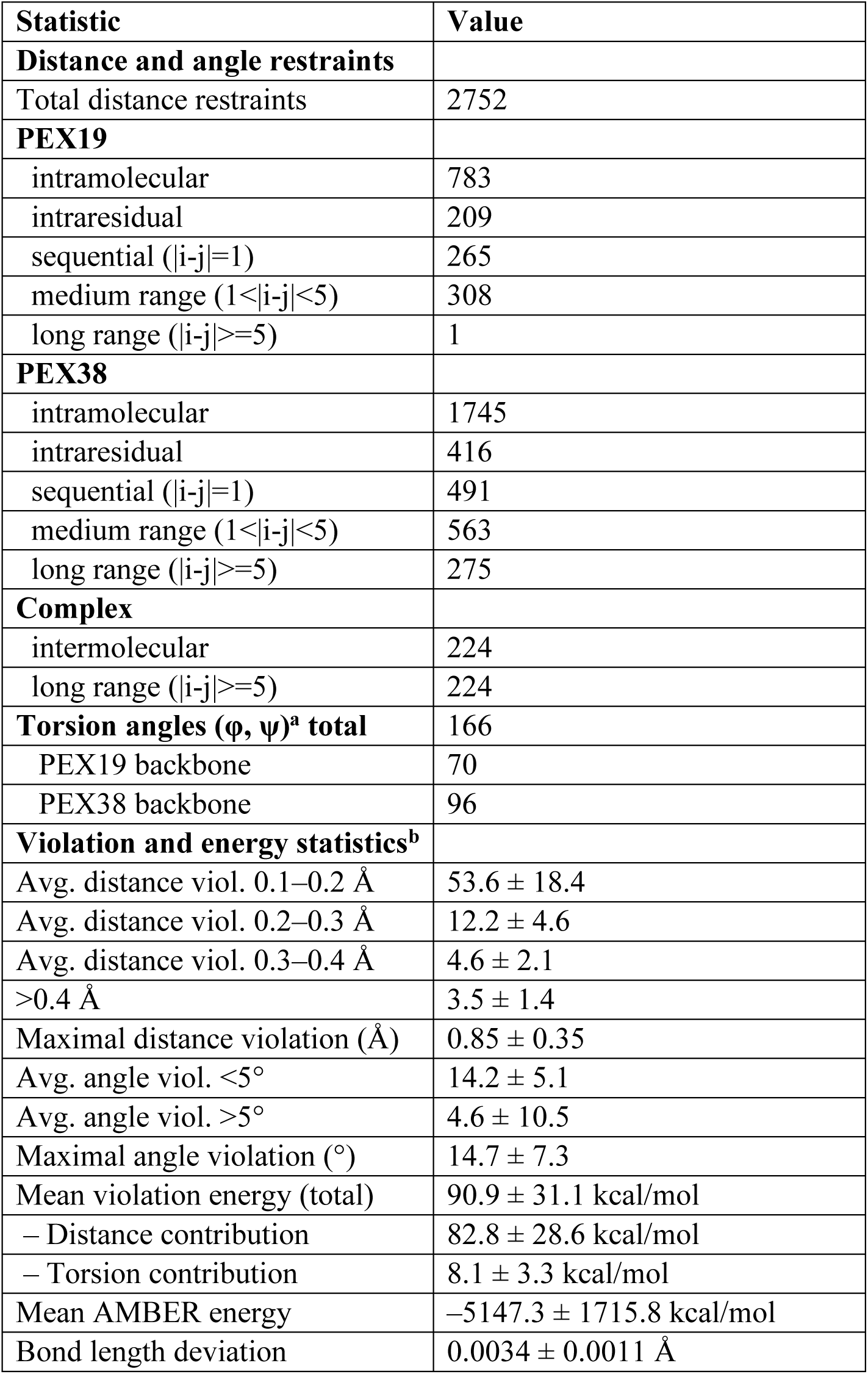
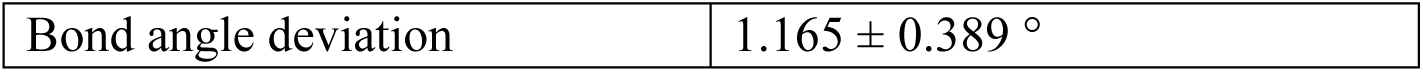
Distance and angle restraints / violation statistics.

**Table S4:** Structural quality (RMSD and Ramachandran)

| Statistic<sup>b,c</sup> | Value |
| --- | --- |
| **RMSD** |  |
| PEX19 backbone atoms | $0.11 \pm 0.05 \text{ \AA}$ |
| PEX19 heavy atoms | $0.50 \pm 0.17 \text{ \AA}$ |
| PEX38 backbone atoms | $0.15 \pm 0.05 \text{ \AA}$ |
| PEX38 heavy atoms | $0.48 \pm 0.09 \text{ \AA}$ |
| All molecules backbone atoms | $0.17 \pm 0.06 \text{ \AA}$ |
| All molecules heavy atoms | $0.51 \pm 0.08 \text{ \AA}$ |
| **Ramachandran<sup>b,c,d</sup>** |  |
| Most favored regions | $95.4 \pm 1.2 \%$ |
| Additionally allowed | $4.6 \pm 1.2 \%$ |
| Generously allowed | $0.0 \%$ |
| Disallowed | $0.0 \%$ |

<sup>a</sup>. Justification for torsion angles, e.g. Molecule 1 TALOS+, Molecule 2 sugar pucker based on homonuclear TOCSY
<sup>b</sup>. Statistics computed for the deposited bundle of 20 violation energy best structure selected out of 30 amber energy best
<sup>c</sup>. Based on structured residue range as defined by cyana command overlay: Molecule 1 (PEX19): 11-20, Chain ID: B (Sequence Range: 1-54); Molecule 2 (PEX38): 100-129, Chain ID: A (Sequence Range: 65-134)
<sup>d</sup>. Ramachandran plot, as defined by the program Procheck<sup>2</sup>

**Table S5:** Strains, Plasmids, and cloning strategy.

| No. | Expression in | Construct | Primer pair | Cloning strategy (Restriction sites) | Cloned in vector |
| --- | --- | --- | --- | --- | --- |
| 1 | *E. coli* | GST-*Tb*PEX19<sub>1-285</sub> (full-length) | RE2926-RE7038 | BamHI/XhoI<br>BamHI/XhoI | pGEX4T2 |
| 2 | | GST-*Tb*PEX19<sub>31-285</sub> (ΔN30) | RE7126-RE7038 | | pGEX4T2 |
| 3 |  | His<sub>6</sub>-*Tb*PEX19<sub>1-285</sub> (full-length) | RE2926-RE7038 |  | pET28a+ |
| 4 |  | GST-*Tb*PEX38 (full-length) | RE7610-RE7611 | EcoRI/XhoI | pGEX4T2 |
| 5 |  | *Tb*PEX19<sub>1-285</sub> (full-length) - His<sub>6</sub> | RE8154-RE8135 | NdeI/XhoI | pET21b |
| 6 |  | *Tb*PEX19<sub>1-50</sub>- His<sub>6</sub> | RE8341-RE8342 | Quick change PCR | pET21b |
| 7 |  | GST-*Tb*PEX3 ΔN44 | 3 |  |  |
| 8 |  | His-SUMO-*Tb*PEX19 1-50-GGY | TbPEX19 SG1+2/3+4 SLIM pETM13S |  |  |
| 9 |  | His-SUMO-*Tb*PEX38 65-134-GGY | TbPEX38 SG1+2/3+4 SLIM pETM13S |  |  |
| 10 | *S. cerevisiae* | GAL4 AD-*Tb*PEX19<sub>1-285</sub> (full-length) | 4 |  |  |
| 11 |  | GAL4 BD-*Tb*PEX38<sub>1-416</sub> (full-length) | RE7508-RE7509 | SalI/NotI | pPC97 |
| 12 |  | GAL4 BD-*Tb*PEX38<sub>69-107</sub> | RE8027-RE8028 |  | pPC97 |
| 13 |  | GAL4 BD-*Tb*PEX38<sub>108-416</sub> | RE8031-RE7509 |  | pPC97 |
| 14 |  | GAL4 BD-*Tb*PEX38<sub>69-133</sub> | RE8027-RE8029 |  | pPC97 |
| 15 |  | GAL4 BD-*Tb*PEX38<sub>134-416</sub> | RE8862-RE8863 | Quick change PCR | pPC97 |
| 16 |  | GAL4 AD-*Sc*PEX5<sub>1-612</sub> | 5 |  |  |
| 17 |  | GAL4 BD-*Sc*PCS60<sub>1-543</sub> | 6 |  |  |
| 18 |  | GAL4 BD-*Tb*PEX38<sub>1-416</sub> (F 73 P) | RE8866-RE8867 | Quick change PCR | pPC97 |
| 19 |  | GAL4 BD-*Tb*PEX38<sub>1-416</sub> (L 96 P) | RE8353-RE8354 |  | pPC97 |
| 20 |  | GAL4 BD-*Tb*PEX38<sub>1-416</sub> (LDL 96-98 PPP) | RE8235-RE8236 |  | pPC97 |
| 21 |  | GAL4 BD-*Tb*PEX38<sub>1-416</sub> (L 96 D) | RE8924-RE8925 |  | pPC97 |
| 22 |  | GAL4 BD-*Tb*PEX38<sub>1-416</sub> (F 73 A) | RE8922-RE8923 |  | pPC97 |
| 23 |  | GAL4 BD-*Tb*PEX38<sub>69-133</sub> (L 96 P) | RE8353-RE8354 |  | pPC97 |
| 24 |  | GAL4 BD-*Tb*PEX38<sub>69-133</sub> (LDL 96-98 PPP) | RE8235-RE8236 |  | pPC97 |
| 25 |  | GAL4 BD-*Tb*PEX38<sub>69-133</sub> (L 96 D) | RE8924-RE8925 |  | pPC97 |
| 26 |  | GAL4 BD-*Tb*PEX38<sub>69-133</sub> (F 73 A) | RE8928-RE8929 |  | pPC97 |
| 27 |  | GAL4 AD-*Tb*Hip<sub>1-388</sub> | RE8112-RE8113 | BglII/NotI | pPC97 |
| 27 |  | GAL4 BD-*Tb*Hip<sub>1-388</sub> | RE8112-RE8113 |  | pPC86 |
| 28 | | GAL4 AD-*Tb*PEX19<sub>31-285</sub> (Δ30aa) | RE7092-RE7095 | Sall/NotI | pPC86 |
| 29 |  | GAL4 AD-*Sc*PEX19<sub>1-342</sub> (full-length) | RE9265-RE9266 (Insert)/ RE9267-RE9268 (Vector) | In-fusion cloning | pPC86 |
| 30 | | GAL4 AD-*Sc*PEX19<sub>31-342</sub> (Δ30aa) | RE9269-RE9270 | Quick change PCR | pPC86 |
| 31 |  | GAL4 AD-*Hs*PEX19<sub>2-299</sub> (full-length) | pAH07<sup>7</sup> |  | pPC86 |
| 32 | | GAL4 AD-*Hs*PEX19<sub>34-299</sub> (Δ33aa) | RE9257-RE9258 | Quick change PCR | pPC86 |
| 33 |  | GAL4 BD-*Hs*SGTA<sub>1-313</sub> | RE9251-RE9252 | Sall/NotI | pPC97 |
| 34 |  | GAL4 BD-*Sc*Sgt2<sub>1-346</sub> | RE9249-RE9250 | Sall/SacI | pPC97 |
| 35 | *T. brucei* | *Tb*PEX38 RNAi (Fragment 1) | RE7528-RE7723 | SpeI/NcoI | Ligate Fragments 1 and 2 |
| 36 |  | *Tb*PEX38 RNAi (Fragment 2) | RE7529-RE7724 | HindIII/NcoI |  |
| 37 |  | *Tb*PEX38 RNAi (Stem-loop) | Ligated fragments 1 and 2 | SpeI/HindIII | p2T7-177 |
| 38 |  | 2×FLAG-*Tb*PEX38; Codon exchanged | Gene synthesis | HindIII/ApaI | pHD1336 |
| 39 |  | 2×FLAG-*Tb*PEX38 (L 96 P); Codon exchanged | RE8858-RE8859 | Quick change PCR | pHD1336 |
| 40 |  | 2×FLAG-*Tb*PEX38 (L 96 D); Codon exchanged | RE8868-RE8869 |  | pHD1336 |
| 41 |  | 2×FLAG-*Tb*PEX38 (LDL 96-98 PPP); Codon exchanged | RE8860-RE8861 |  | pHD1336 |
| 42 |  | 2×FLAG-*Tb*PEX38 (F 73 P); Codon exchanged | RE8864-RE8865 |  | pHD1336 |
| 43 |  | GFP-*Tb*PEX38 (full-length) | RE7573-RE7575 | BstBI/ApaI | pGC1 |
| 44 |  | *Tb*PEX38-GFP (full-length) | RE7573-RE7574 |  | pGN1 |
| 45 |  | GFP-SKL | 8 |  |  |

**Table S6:**
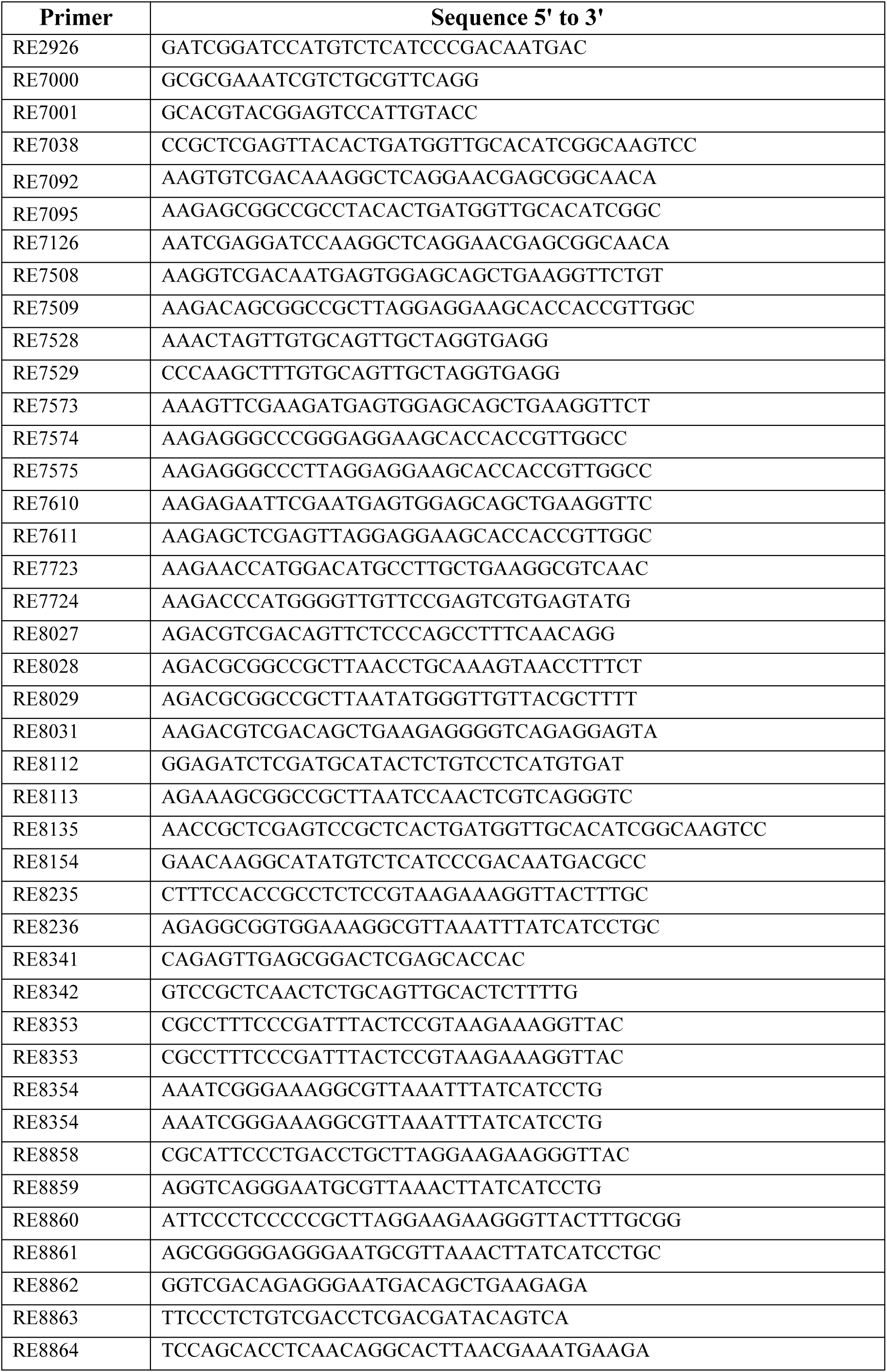
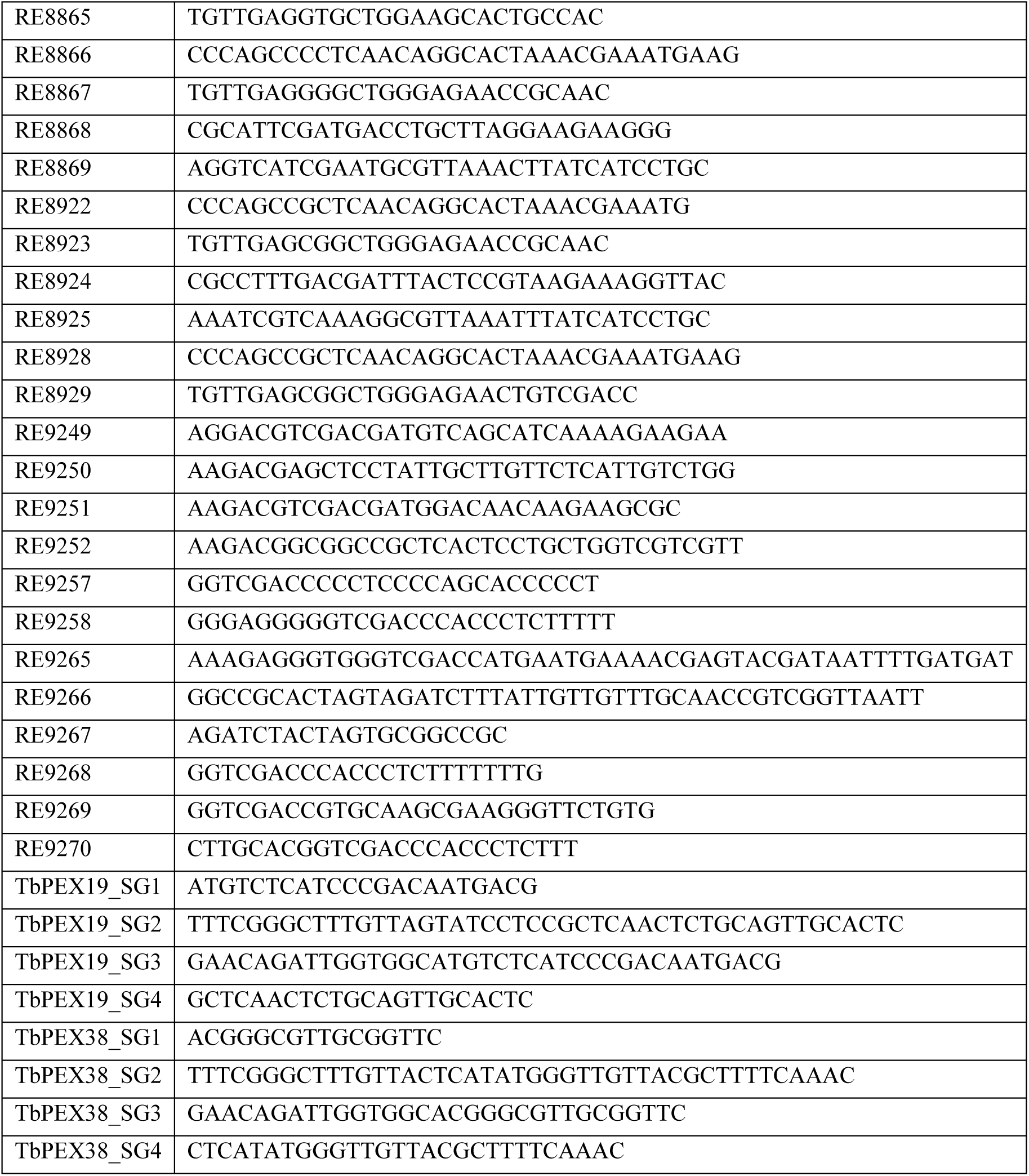
Oligonucleotides.

